# Scalable covariance-based connectivity inference for synchronous neuronal networks

**DOI:** 10.1101/2023.06.17.545399

**Authors:** Taehoon Kim, Dexiong Chen, Philipp Hornauer, Sreedhar Saseendran Kumar, Manuel Schröter, Karsten Borgwardt, Andreas Hierlemann

## Abstract

We present a novel method for inferring connectivity from large-scale neuronal networks with synchronous activity. Our approach leverages Dynamic Differential Covariance to address the associated computational challenges. First, we analyze spike trains generated from Leaky Integrate-and-Fire network simulations and evaluate the performance of several off-the-shelf multivariate connectivity inference methods. Next, we introduce a new approach, Fractional Dynamic Differential Covariance (FDDC), and demonstrate that it consistently outperforms the other methods. Finally, we apply FDDC to experimental data to assess the topological organization of inferred graphs of in vitro neural network recordings obtained using high-density microelectrode arrays (HD-MEAs). Our results indicate that FDDC-derived graphs exhibit a significant negative correlation between small-worldness and measures of network synchrony. In contrast, graphs inferred through the well-established pairwise correlation method do not show such a correlation. This finding implies that the graphs obtained through FDDC provide stronger evidence in support of the theoretical notion that networks with clustered connections tend to exhibit higher levels of synchronizability. We hypothesize that our findings have implications for the development of scalable connectivity inference methods for large-scale neural network data.

## Introduction

Large-scale electrophysiological methodology, such as high-density microelectrode arrays (HD-MEAs) play an important role in studying neuronal activity and for gaining insights into the functioning of neural networks (1). Recent years have seen significant progress in this field, particularly an increase in the number of recording electrodes, resulting in improved spatio-temporal recording resolution (2–5). However, such gains in information content of neuronal recordings come with a significant increase in data volume - which can pose several computational challenges. The sheer volume of data generated by these arrays can quickly become overwhelming, making it difficult to process the data in a timely manner. These challenges become even more pronounced when inferring functional connectivity from the recordings, because an increase in the number of recording electrodes often corresponds to an increase in the number of recorded neurons. As connectivity inference methods often rely on pairwise calculations (1), the inference becomes computationally intensive as the number of recorded neurons grows. Moreover, many of today’s inference methods rely on the assumption that neurons in the network exhibit asynchronous firing activity, making them less suitable for inferring connections during synchronous network activity (6–9). Although approaches to filter out time windows of synchronized activity exist, such as detecting population bursts (10) or factoring out common activity through multivariate modeling (11, 12), they introduce additional computational load and conceptual complications. Therefore, previous studies have investigated multivariate methods to uncover the true synaptic connectivity between neurons by mitigating the influence of signals from other neurons in the network. Baker et al. (13) examined the theoretical feasibility of recovering neural connectivity and found that utilizing the precision matrix, derived from the inverse of the covariance matrix of spike trains, seems to be an effective approach for inferring underlying neuronal connections across diverse experimental scenarios. Furthermore, there is a considerable body of literature on connectivity inference from the perspective of maximum likelihood estimation with sparsity assumptions, often emphasizing the sparse estimation of precision matrices. While the majority of such approaches has focused on inferring functional connectivity between brain regions (14–20), relatively few studies have applied the latter to the inference of connectivity between individual neurons (21, 22). Although these methods hold considerable promise for scaling up to large networks (23), further exploration is required to evaluate their efficacy for large-scale neuronal networks (e.g., n>1000 neurons) and to assess their potential limitations.

The main goal of this study was to tackle the computational challenges associated with inferring connectivity in largescale, synchronous neural networks. We, therefore, applied and evaluated scalable methods on simulated data and proposed an improved connectivity inference method. Specifically, we compared the performance of off-the-shelf multivariate connectivity inference methods that rely on the covariance computed from the spike trains of observed neurons in a network. We focused on these methods as more sophisticated methods may not scale for networks with large numbers of neurons (i.e. n>1000 neurons).

As a testing ground, we used spike trains generated from random networks with 2000 Leaky Integrate-and-Fire (LIF) neurons that mimicked synchronous network firing activity of hippocampal networks through adaptation of synaptic currents(24). We compared the performance of these inference methods based on their AUROC (Area Under the Receiver Operating Characteristic Curve) and AUPRC (Area Under the Precision-Recall Curve) values. Then we made two modifications to the best method, Dynamic differential covariance (DDC)(25), and introduced an improved method, Fractional Dynamic Differential Covariance (FDDC), which consistently showed superior performance. Finally, we applied FDDC to infer connections from in vitro neural recordings obtained using high-density microelectrode arrays (HD-MEAs) and compared the resulting graphs to those obtained from a well-studied pairwise inference method (cross correlogram (CCG)). Analysis of the topology of the inferred networks revealed that CCG-derived graphs showed higher small-worldness compared to the FDDC-derived graphs. Moreover, FDDC-derived graphs showed a clear negative correlation between small-worldness and a measure of network synchrony (Participation ratio) across different graph densities, while CCG-derived graphs did not. These results highlight the utility of fractional differentiation and inversion of the covariance matrix for extracting pairwise relations between neurons in synchronous networks. We argue that our findings have important implications for developing connectivity inference methods that can handle networks with large numbers of neurons.

## Materials and Methods

In this section, we outline the LIF simulation that was used to simulate the activity of neural networks. Then we provide an overview of the different connectivity inference methods and describe the in vitro neural recordings used in this study. Lastly, we address how synchrony in the experimental neural recordings was quantified, as well as how we evaluated the topologically clustered organization of the inferred neuronal connectivity.

### Simulating network activity with an adaptive exponential integrate-and-fire model

We employed an adaptive exponential integrate-and-fire model, as described in (26), for simulating network activity. Our implementation was based on the methodologies presented in (24, 27). In this context, we denote *C* the membrane capacitance and *V*_*i*_ the membrane potential of neuron *i*. Additionally, *E*_*L*_ refers to the leaky membrane potential and *g*_*L*_ is the leakage conductance.

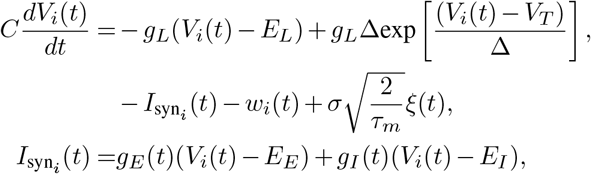

where *V*_*T*_ represents the effective threshold for generating spikes. When the membrane potential *V*_*i*_ reaches *V*_*T*_, it is reset to *V*_reset_ and remains constant for the duration of *T*_refrac_ (see Table S1). The threshold slope factor is denoted by ∆, and 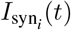 (*t*) is the total synaptic current received by neuron *i* from its presynaptic neurons. The reversal potential for excitatory and inhibitory synapses is expressed as *E*_*E*_ and *E*_*I*_, respectively. *ξ*(*t*) is a Gaussian white noise with *σ* being the noise strength. Lastly, *τ*_*m*_ refers to the time constant of the noise, which can be calculated as *τ*_*m*_ = *C/g*_*L*_.

The following equation represents the change in excitatory and inhibitory conductance, denoted *g*_*E*_ and *g*_*I*_ respectively:

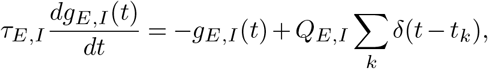

Excitatory/inhibitory conductance (*g*_*E*_/*g*_*I*_) increases by a quantity of *Q*_*E*_/*Q*_*I*_ each time the presynaptic excitatory/inhibitory neuron fires at a specific time *t*_*k*_. Furthermore, the synaptic conductance *g*_*E*_/*g*_*I*_ undergoes an exponential decay, following a corresponding time constant *τ*_*E*_/*τ*_*I*_.

The following equation describes the adaptation current of neuron *i*, represented by *w*_*i*_:

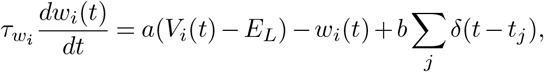

where the parameter *a* models the sub-threshold adaptation. When neuron *i* fires at time *j, w*_*i*_ increases by a value of *b*. Additionally, *w*_*i*_ decays exponentially with the time constant 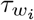.

All LIF network simulations consisted of n = 2000 neurons with 80% excitatory and 20% inhibitory neurons. All inhibitory neurons were set to show no adaptation, e.g., (*a, b*) = (0,0), while excitatory neurons were set to specific (*a, b*) parameters (i.e., obtained through parameter search) to implement adaptation. All neurons were randomly connected with a connection probability of 2%. For each connected edge, the synaptic delay was implemented by randomly selecting a value in the range of [0 5] ms. At the beginning of the simulation, a shared Poissonian spike train with the same excitatory strength *Q*_*E*_ and a mean firing rate of 500 Hz was applied to drive all neurons in the network for 100 ms. Afterward, the network was purely driven by its own recurrent activity. For the downstream analysis, the first second (s) of activity was removed to consider only spontaneous recurrent activity. All described equations were numerically integrated with a step size of 0.1 ms using the ‘Brian2’ python library^1^. Network activities were simulated for 20 min as longer simulations could not be handled with the computing resources utilized in this study (e.g., computing instances with 64 GB RAM). All parameters used for the simulation are provided in the Supplementary material (Table S1).

### Parameter search

A parameter search was performed to select the set of parameters (*a, b, g*_*E*_, *g*_*I*_) for generating network activities ranging from asynchronous irregular firing activity (Async) to synchronous firing activity (Sync). The search was performed in two steps. First, by fixing the adaptation parameters (*a, b*) = (1*µ*S, 5nA) to implement “weakly adapting cells” as stated in (24), excitatory/inhibitory conductances (*g*_*E*_, *g*_*I*_) were searched in the range of [0 100] with a step size of 5 nS (400 combinations)(Fig. S1). For each parameter set, the network activity was simulated for 5 s. We selected (*g*_*E*_, *g*_*I*_) = (10, 65) as the default excitatory/inhibitory conductances, as this set of parameters showed spontaneous firing activity with the highest excitatory conductance (*g*_*E*_) without saturation in the network firing rate (e.g., 200 Hz for *T*_refrac_ = 5 ms)(Fig. S1). As the second step, (*a, b*) was searched in the range of [0 80] with a step size of 1 *µ*S, nA) (6400 combinations). 5 s of activity was simulated for each parameter set, and the simulated activities were quantified in how synchronous the spike trains were in the network.

### Characterization of network activity

To characterize the network firing synchrony of LIF simulations, we computed the mean coefficient of the variation of interspike intervals (CV_*ISI*_) and mean pairwise correlation (MPC) as implemented in Destexhe (2009) (24). We only considered neurons showing firing rates *>* 0.01 Hz for the following quantities. CV_*ISI*_ measures the temporal regularity of the network firing activity and is calculated as follows.

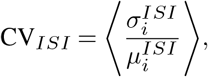

where 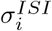 and 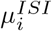 denote standard deviation and mean of the ISIs of neuron *i*. The bracket 〈〉 indicates an average over all neurons in the network.

The MPC, on the other hand, measures the degree of synchrony between all neurons in a network.

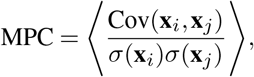

where **x**_*i*_, **x**_*j*_ denote binned spike trains of neurons *i* and *j* computed using a bin size of 5 ms. We considered the simulated network activity to show asynchronous activity only if CV_*ISI*_ *>* 1 and MPC *<* 0.1. If a simulated network did not satisfy one of these criteria, the network was considered to exhibit synchronous firing activity (24).

### Parameter selection for LIF network simulations

The parameter search for the LIF networks showed that with *b >* 60 and *a < b*, simulations were more likely to show synchronous network activity (Fig. S1). This observation was in agreement with previously reported values (24, 28). From all 6400 combinations of (*a, b*), the parameter sets that generated simulations without lasting spontaneous activity or with very high network firing rates (30 Hz) were discarded. For both asynchronous and synchronous networks, we chose two parameter sets that exhibited the lowest or highest MPC, respectively. We then generated five 20 min simulations for each parameter set. From these ten network simulations of each network type, we selected the five most synchronous or asynchronous networks and used them for the inference tasks.

### Perturbation of LIF network simulations using white noise

The inference was performed on noise-free LIF networks unless specified otherwise. For noise-perturbed simulations, the same connections and synaptic delays as in the non-perturbed counterpart were used. A wide range of noise currents was applied with values both smaller and larger than that of the leaky current (e.g., *g*_*L*_(*V*_*i*_(*t*) − *E*_*L*_)), as we observed qualitative changes in network behavior in terms of MPC and firing irregularity (CV_*ISI*_) in cases where the noise was larger than the leaky current (Fig. S2).

### Functional connectivity inference methods

In this study, we focused on connectivity inference methods that could scale to large-scale networks (n *>* 1000 neurons). When binning of spike trains was necessary, we used 5 ms as a bin size to reflect the synaptic delay of the simulation unless specified otherwise. We first implemented a well-established pairwise inference method that is widely used to study functional connectivity of neural circuits, the results of which served as a baseline in this study (1, 29). We anticipated that the method would perform exceptionally well for asynchronous networks. However, we expected its performance to be suboptimal for synchronous networks, where the spike time correlations between neurons would be masked by the synchronous firing activity of the entire network. We then assessed how the selected set of scalable multivariate inference methods compared to the pairwise method and drew conclusions on their usefulness.

### Pairwise cross-correlograms

To assess pairwise relations between neurons, we adapted the cross-correlogram method (CCG) implemented in English et al. (30). For all neuron pairs in the network, cross-correlograms were computed using a bin size of 1 ms for a time window [-50 +50] ms. Then the count in each bin was compared to the baseline value (*λ*_base_), which was generated as follows. The observed crosscorrelograms were convolved with the partially hollow Gaussian Kernel (31) with a standard deviation of 10 ms, with a hollow fraction of 60% to model baseline (30). We estimated the probability of observing a spike count *x ≥ n* in the postsynaptic bin *t* using the Poisson distribution with a continuity correction (31).

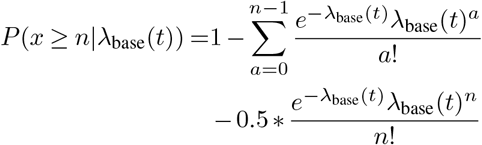

We also computed 1 *− P* (*x≥ n* |*λ*_base_(*t*)) to estimate the likelihood that the connection was inhibitory. For each neuron pair, we took the maximum (peak, excitatory connection) and the minimum (trough, inhibitory connection) in the postsynaptic bins [0 5] ms, estimated the probability of respective values and compared the negative log-likelihoods. If the negative log-likelihood was larger for the maximum/minimum, we considered the edge to be an excitatory/inhibitory connection. As a result, we obtained a directed, weighted connectivity with each edge representing the negative log-likelihood.

### Network deconvolution

We implemented Network deconvolution (ND) (32) to test if synaptic connectivity can be recovered from observed covariance matrices containing direct and indirect interactions between neurons. We binned the spike trains of the neurons in the network with the predefined bin sizes (5 ms) to match the LIF simulations. Then, these binned spikes counts were used to compute the covariance matrix (**S**), which represents the variance between the firing activity of neurons given the timescale (bin size). Under the strong assumption that the covariance matrix is a result of the graph signaling process of the latent graph **A**, and the graph signal propagation is linear, time-invariant, and flow-preserving, the relation between the latent graph **A** and the covariance matrix **S** can be written as follows (32).

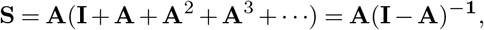

which equivalently states **A** = **S**(**I** + **S**)^*−***1**^. From this relation, we computed ND in three steps. First, **S** was rescaled to ensure that the largest absolute eigenvalue of the **A** was strictly smaller than one. This was done by choosing a linear scaling factor *α* (**S**_*scaled*_ = *α***S**):

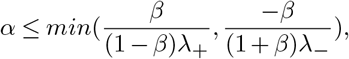

where *β* is the target value for the largest absolute eigenvalue of **A** (*β <* 1), and *λ*_+_/*λ* is the largest/smallest eigenvalue of **S**. As a second step, the **S**_*scaled*_ was decomposed to get eigenvalues and eigenvectors such that **S**_*scaled*_ = **UΣ**_**S**_**U**^*−***1**^. As the final step, we computed the diagonal eigenvalue matrix of **A, Σ**, using the relation 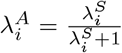 Here, 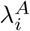 was the *i*-th component of **Σ**_**A**_ and 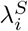 was the *i*-th component of **Σ**_**S**_. The latent graph **A** was then computed as **A** = **UΣ**_**A**_**U**^*−***1**^, and we considered **A** as the estimate of the connectivity. In this study, we chose *β* = 0.99 to get a sparse estimate of connectivity. For more details on the ND approach, we refer the readers to Feizi et al. (32).

### Graphical lasso

The covariance matrix, derived from binned spike trains (bin size = 5 ms), can be inverted to obtain a precision matrix. Each entry in the precision matrix, **P**_*ij*_, quantifies the correlation between neuron *i* and *j* while factoring out all the activity of other neurons in the network. This is particularly valuable for identifying true pairwise correlations between neurons if the network exhibits synchronous firing activity. Instead of directly inverting the covariance matrix, the precision matrix can be estimated using sparse regularization methods, resulting in a sparser estimate of the precision matrix. This approach is beneficial when the covariance matrix is rank-deficient or too large for calculating the inverse. A widely used method among them is the graphical least absolute shrinkage and selection operator (graphical LASSO (Glasso))(33), which is defined as follows.

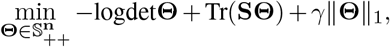

where 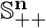 denotes a positive-semidefinite (PSD) matrix of dimensions *n× n*. **S** is the observed covariance matrix, and **Θ** is the estimated precision matrix. ‖.‖_1_ is an entrywise *l*_1_-norm, and *γ* is regularization penalty. This objective can be optimized using the alternating direction method of multipliers (ADMM) (34) with an augmented Lagrangian, given by

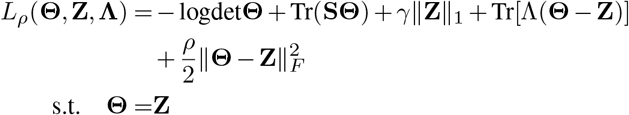

where **Λ** is a dual variable and *ρ* is a penalty parameter (*ρ >* 0). ‖·‖ _*F*_ is a Frobenius norm, e.g., a square root of the sum of squares of entries in a matrix. By additionally setting a scaled dual variable **U** = **Λ***/ρ*, we get simplified update rules.

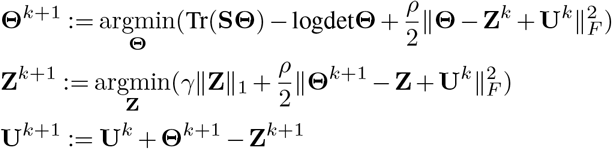

where Θ is updated by setting the gradient to zero.

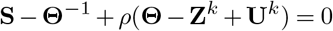

By rewriting the equation,

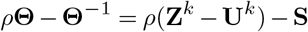

we can separate Θ on the left-hand side of the equation. The right-hand side of the equation can be decomposed as

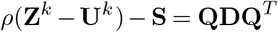

where **D** is the diagonal matrix of eigenvalues. If we consider 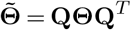, we find the 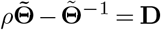 and arrive at the following solution for each diagonal element of 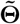.

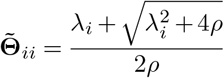

We update **Θ**^*k*+1^ by setting 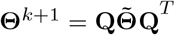. From (35), **Z** update is simplified to the soft thresholding operator *F* as follows.

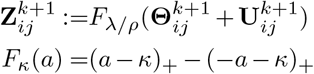

The ADMM was implemented using ‘gglasso’ Python library^2^ (36). We computed eight logarithmically spaced *γ* values in the range [10^*−*1.5^, 10^0.5^] (i.e., ‘numpy.logspace(0.5, -1.5, 8)’). The best regularization parameter *γ* was selected using the extended Bayesian information criterion (eBIC) as suggested in (37). Other than *γ*, we used default parameters for both ADMM and eBIC computation unless specified otherwise.

### Differential covariance

We implemented Differential covariance (Dcov) (38) to investigate the utility of directed temporal covariance in inferring synaptic connectivity. We represent the column vector **x**(*t*) that has the *i*-th entry equal to the z-transformed number of spikes or firing rate of the neuron *i* at time *t*. Dcov is defined as the covariance between the **x**(*t*) and the derivative.

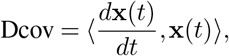

where ⟨, ⟩ denotes time averaged vector outer product and differentiation is computed as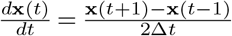The differentiation term in Dcov makes it a directed measure of connectivity.

### Dynamic differential covarianceb

Dynamic differential covariance (DDC)(25) can be estimated as a matrix product between the Dcov and the precision matrix (i.e., the inverse of the covariance matrix).

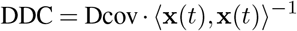

DDC improves upon Dcov by factoring out the influence of common input sources. For further proofs and characteristics of DDC, we refer readers to Chen et al. (25).

### Fractional dynamic differential covariance

If a time series exhibits long decay in autocorrelation, taking fractional (non-integer) differentiation can preserve the lasting trend in the time series which would otherwise be lost for full integer differentiation (39). Fractional differentiation with a fractional order *β*, is defined as:

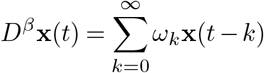

*where ω0 = 1, ω1 = −β*, and 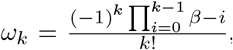 for *k ≥*2. If *β* = 1, we retrieve full-integer differentiation. In this study, we implemented up to 10-th order (*k* = 10) to compute fractional differentiation using Python library ‘fdiff’ ^3^. By substituting the differentiation in DDC with fractional differentiation, we could compute the Fractional dynamic differential covariance (FDDC).

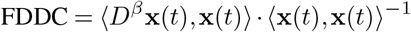

### Spike train convolution

One strategy to increase the number of sample points from spike trains is to compute instantaneous rates (40). Instantaneous rates can be computed by convolving the spike trains using predefined kernels. We used two commonly used functions to generate instantaneous rates: alpha-function kernels (Alpha kernels) and Gaussian kernels. The Gaussian kernel has the form:

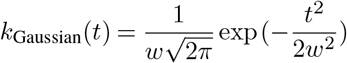

We denote a filter size (duration in ms) as *w*. As an asymmetric counterpart, we applied Alpha kernels as a complementary approach. The Alpha kernel is a decaying exponential function with a characteristic time constant (*τ*), which facilitates the modeling of postsynaptic currents (41). The kernel was defined as:

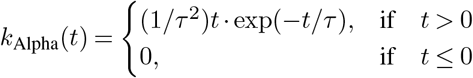

We considered 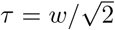 for the consistency in notation with the Gaussian kernel. For both kernels, filter sizes (*w*) of 1, 3 ms were used to generate instantaneous rates. The filter size of 1 ms corresponded to the typical duration of an action potential (i.e., *<* 5 ms), and 3 ms corresponded to the timescale of longer synaptic delays (i.e., *<* 20 ms) in experimental studies (42, 43). We sampled points at a frequency of 1 kHz from these instantaneous rates.

### Evaluation of inference performance

We assessed the performance of each connectivity inference method against the ground truth by calculating the Area Under the Receiver Operating Characteristic Curve (AUROC) and the Area Under the Precision-Recall Curve (AUPRC). The absolute values of computed inference scores (weights) were used to compute AUROC and AUPRC. Additionally, we investigated the impact of simulation duration on inference performance by testing 5, 10, 15, and 20 min of simulated activity. To determine whether multivariate methods benefited from an increase in the number of observed neurons, we sampled 500, 1000, and 1500 neurons for the analysis. We randomly sampled three times from each simulated network, so for one condition (e.g., 5 min simulation length, 500 sampled neurons), we reported the mean and standard deviation of AU-ROC/AUPRC values from 15 instances (i.e., five network simulations *×* three samples). We also compared wall-clock time to assess the scalability of the inference methods. While all multivariate methods were implemented in Python, CCG was implemented using C and Matlab adapted from English et al. (30)(see Section “Data and code availability”). Python implementation of CCG using a standard multiprocessing library (‘map’) was roughly five times slower than the values reported in this study. The computation time was measured excluding the computation for preprocessing (e.g., binning, kernel convolution of spike trains). For all inferences, we used computing instances with the same specification with 8 CPU cores (Intel(R) Xeon(R) Gold 6154 CPU @ 3.00GHz) and 64 GB of RAM.

### Recording neural network activity in vitro

#### High-density microelectrode array recordings

HD-MEA recordings used in this study were part of the dataset reported in a prior study (44). We performed recordings with commercially available 6-well high-density microelectrode array (HD-MEA) plates (MaxTwo system by Maxwell Biosystems, Zurich, Switzerland). Each well included a CMOS-based HD-MEA(2) featuring 26,400 electrodes arranged in a 120 × 220 electrode grid with a microelectrode center-to-center spacing (pitch) of 17.5 *µ*m. The overall sensing area of this HD-MEA was 3.85 × 2.10 mm^2^. The HD-MEA enabled the simultaneous recording of up to 1020 electrodes at a sampling rate of 10 kHz. Recordings were performed in an incubator chamber of the MaxTwo system at 37°C and 5% CO_2_/ 95% O_2_ and were conducted at days in vitro (DIVs) 22–25. Each recording started with a whole-array ‘activity scan’ to determine the active electrodes on the HD-MEA. The activity scan consisted of 29 dense electrode configurations to scan through the entire sensing area of the electrode array; each configuration was sequentially recorded for 60 s. From the activity scan, up to 1020 electrodes were selected by prioritizing electrodes with high firing rates. Using the selected electrodes, we recorded two hours of spontaneous activity from 22 HD-MEA wells or neuronal cultures.

#### Hippocampal dissociated neuronal culture

Rat primary neurons were acquired from the dissociated hippocampus of Wistar rats on embryonic day (E) 18, using the protocol described in (45). All animal experiments were conducted in accordance with the approved guidelines and Swiss federal laws on animal welfare, with approval from the Basel-Stadt veterinary office. HD-MEA wells were sterilized in 70% ethanol for 30 min prior to cell plating. Afterwards, the ethanol was removed, and the wells were rinsed three times with sterile distilled water before being left to dry. The HD-MEA wells were then coated with a layer of 0.05% polyethylenimine (Sigma-Aldrich, Buchs, Switzerland) in a borate buffer (Thermo Fisher Scientific, Waltham, MA, United States) to render the surface more hydrophilic. Then, a thin layer of laminin (Sigma-Aldrich, 0.02 mg/mL) in Neurobasal medium (Gibco, Thermo Fisher Scientific) was applied to the array and incubated for 30 min at 37°C to promote cell adhesion. We dissociated hippocampi of E18 Wistar rat in trypsin with 0.25% EDTA (Gibco), followed by trituration. Cell suspensions of 15, 000 cells in 7 *µ*L were then plated on top of the electrode arrays. The plated wells were incubated at 37°C for 30 min before adding 2 mL of the plating medium. The plating medium consisted of Neurobasal, supplemented with 10% horse serum (HyClone, Thermo Fisher Scientific), 0.5 mM Glutamax (Invitrogen, Thermo Fisher Scientific), and 2% B-27 (Invitrogen). After 5 days, 50% of the plating medium was replaced with a growth medium, containing Brainphys medium supplemented with SM1 and N2-A (Stemcell technologies, Cologne, Germany). For the rest of the experiments, medium changes were performed twice a week using the same Brainphys-based medium. The wells were kept inside a humidified incubator at 37°C and 5% CO_2_/ 95% O_2_.

#### Spike-sorting and quality control of sorted units

For each HD-MEA/well, the recordings were filtered, and spike-sorted using ‘Kilosort2’ (46); the applied parameters are stated in Table S2. To be included in subsequent analyses, all inferred spike-sorted units had to pass a quality control: First, we removed units with a firing rate below 0.05 Hz and higher than 30 Hz. Then we computed the refractory period violation ratio, which was calculated as the fraction of interspike intervals (ISIs) less than 2 ms (47). In practice, we quantified the number of spikes within the [*±*2 ms] bins of the spike train autocorrelogram (ACG) and then computed the fraction between this count and the total number of spikes in a larger range of the ACG [*±* 50 ms]. Any template exceeding a refractory period violation ratio of 0.3 was removed. Based on these preprocessing steps, the obtained units were considered to originate from single neurons.

#### Characterizing the spontaneous activity of in vitro hippocampal networks

##### Participation ratio

We used the Participation Ratio (PR) to measure the correlated firing activity in each network recording, normalized by the number of neurons. The calculation and interpretation of the PR were based on the implementation of Recanatesi et al.(48). First, the spike trains of the baseline recordings were binned using a window size of 5 ms. The resulting binned spike trains were z-transformed and used to compute inner products, generating a correlation matrix. From the eigendecomposition of the correlation matrix, we collected the eigenvalues to assess the level of correlated activity between neurons. The participation ratio was defined as

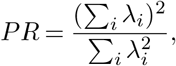

where *λ*_*i*_ is *i*-th eigenvalue of the correlation matrix. The resulting PR value indicates the number of principal components that were necessary to explain 80% to 90% of the total variance for typical Principal component analysis (PCA) eigenspectra (49). We then normalized the PR, by dividing it by the number of neurons (*N*) in the network.

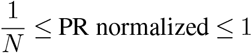

A normalized PR value of *<* 0.8, suggests that the majority of the variance in the network activity could be explained by less than 80% of principal components (48–50).

##### Small-worldness of inferred networks

The small-worldness of a network measures the extent to which a network exhibits both high local clustering and short average path lengths between nodes (51). To conclude whether a network shows small-worldness, Downes et al. (52) defined Small-World Index (SWI) as a ratio between clustering coefficient (CC) and characteristic path length (L_path_). CC was defined as the fraction of possible triangles in the network that are present, while the L_*path*_ was the average number of steps along the shortest paths for all possible pairs of network nodes (51). In practice, CC and L_path_ need to be normalized against the expected values computed from random networks with the same number of nodes and edges as the original graph (29). If SWI *>* 1, the network is considered to show small-worldness. To investigate the differences in the inferred network structure from in vitro recordings, we inferred connectivity from the recordings using both FDDC and CCG. For each case, we applied thresholds to generate multiple binary, directed graphs with density (*ρ*) of 5, 10, 15, and 20%. CC and the L_path_ were then computed from each thresholded graph, and lattice surrogates (*n* = 30) were generated by rewiring the edges in the thresholded graphs while preserving the distribution of in-degree and out-degree across nodes (53). From lattice surrogates, the average CC and L_path_ were calculated. Then SWI was computed as the ratio between the normalized CC and L_path_.

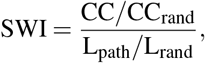

where CC_rand_ and L_rand_ were the average clustering coefficient and characteristic path length of the lattice surrogates. The generation of lattice surrogate networks and subsequent computation of SWI was implemented using the Python library ‘bctpy’ ^4^.

##### Measurement of autocorrelation in convolved spike trains

To assess the temporal dependency present in a sequence of convolved spike trains, we computed the normalized autocorrelation.

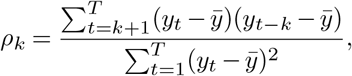

where *y*_*t*_ is the value of the convolved spike train at time *t*, and 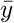 is the mean of the convolved spike train. *T* is the total number of observations and *k* is the lag. In this study, *T* was set to 20 min with the bin size of 1 ms, and *k* was computed up to 20 (20 ms). After computing *ρ*_*k*_, we normalized the resulting values by dividing them by the maximum value *ρ*_0_ (*k* = 0). This ensured that the normalized autocorrelation values ranged from -1 to 1. In the case of simulated networks, we randomly sampled 500 neurons and computed the autocorrelation only for those neurons that had a firing rate higher than 0.1 Hz. For the experimental data, we also computed the autocorrelation only for neurons that had a firing rate higher than 0.1 Hz. We characterized the degree of structure in the spike trains by computing the average area under the curve (AUC) of the autocorrelation curves. These AUC values were compared with the AUC values of randomized surrogates, which shuffled spike times randomly while keeping the total number of spikes.

## Results

In this section, we first characterize the simulated networks and pinpoint the condition under which the pairwise inference approach falls short. After identifying the failure mode, we assess the performance of the multivariate inference methods. Then, we build up on the most promising method and apply a new inference approach, FDDC. Lastly, we investigate the FDDC further by applying it to HD-MEA network recordings from in vitro neuronal cultures and compare the inferred FDDC-derived connectivity graphs with those derived with the CCG method.

### Simulated networks with asynchronous and synchronous network activity

The simulated asynchronous and synchronous networks showed clear differences in their average population firing rates and in both measures of network synchrony (MPC, CV_*ISI*_). The asynchronous networks showed average population firing rates of 31 Hz (standard deviation 0.7 Hz), and the synchronous networks exhibited average population firing rates of 20 Hz (1.3 Hz). The average CV_*ISI*_ for the asynchronous networks was 1.7 (0.04), and 1.1 (0.11) for the synchronous networks. Moreover, the asynchronous networks showed on average lower MPC values (0.02 (0.002)) compared to the MPC values obtained for the synchronous networks (0.24 (0.06); Fig. 1).

**Fig. 1.**
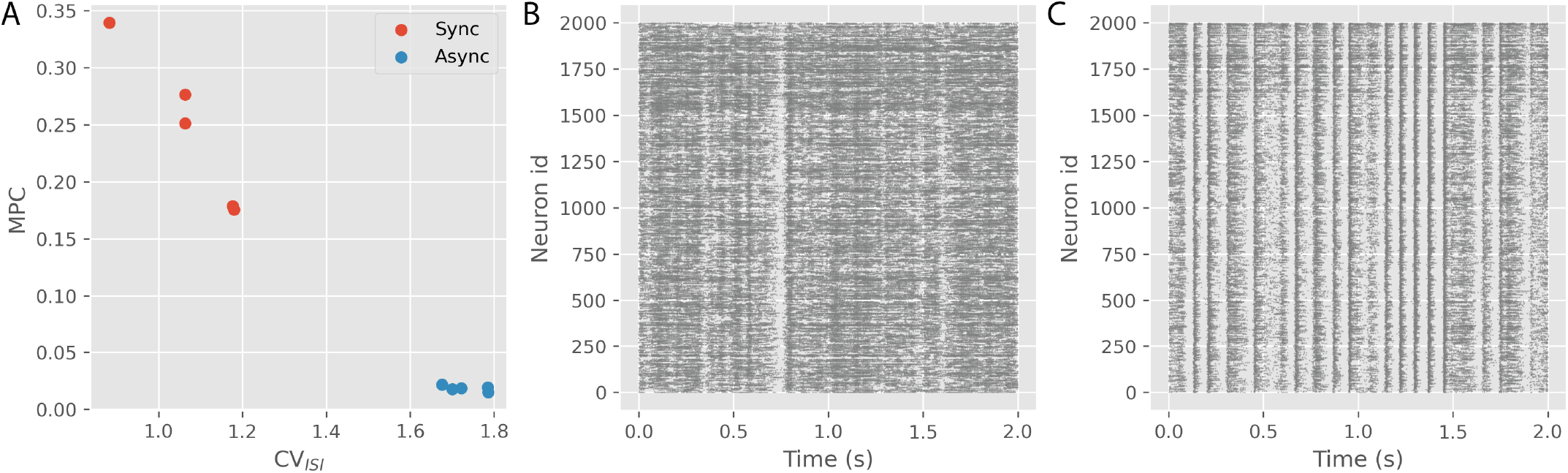
LIF network simulations. (A): There was a clear difference in the measures of network synchrony (mean pairwise correlation (MPC), CV_*ISI*_) between the two network simulation types. Asynchronous networks showed average population firing rates of 31 Hz (standard deviation 0.7 Hz), while synchronous networks exhibited average population firing rates of 20 Hz (1.3 Hz). (B): A spike raster plot showing 2 seconds of the network firing activity from an asynchronous network. (C): A raster plot showing 2 seconds of network firing activity from a synchronous network. In contrast to the asynchronous network (depicted in Panel B), a more prominent synchronized firing pattern is observed.

### Pairwise inference using cross-correlograms fails to recover connectivity of simulated synchronous network activity

Upon examining the inference methods on both asynchronous and synchronous simulated networks, we discovered that the CCG performed exceptionally well for the asynchronous networks while only attaining performance close to random success for the synchronous networks (i.e., AUROC *≈* 0.5, AUPRC *≈* 0.02). An overview of the performance across all methods is provided in Table 1 (For more details, see also Table S3 and Fig. 2A). Although none of the multivariate methods outperformed CCG for the asynchronous networks, DDC showed the best performance for the synchronous networks (Table 1, S4, S5). Moreover, as the number of observed neurons (n) increased, the average AUROC and AUPRC scores for DDC also showed improvement (Fig. 2B). In contrast, Glasso demonstrated inference performance that was close to random estimations, and ND, Dcov showed marginally better performance compared to Glasso (Table S4, S5, Fig. S3, S4). We additionally evaluated the AUROC and AUPRC values for undirected methods, such as ND and Glasso, by transforming the directed ground truth connectivity into undirected connections. The resulting scores remained consistently low for both methods (Table S6).

**Table 1.** Comparison of inference performance. The performance of each inference method for the simulated network activities, based on 1500 sampled neurons and 20 min of simulation, is presented in the table. The average values rounded for two significant digits are reported for the computation time. For the asynchronous networks, CCG showed the best performance with the highest AUROC and AUPRC, but failed to recover the ground truth connections for the synchronous networks. DDC showed the best performance for the synchronous networks in addition to faster computation.

| Network act. | Method | AUROC | AUPRC | Comp. time (s) (averaged) |
| --- | --- | --- | --- | --- |
| Async | **CCG** | **$0.93 \pm 1.1\text{e-}2$** | **$0.55 \pm 2.4\text{e-}2$** | **3300** |
| | ND | $0.63 \pm 5.4\text{e-}3$ | $0.06 \pm 2.4\text{e-}3$ | 9 |
| | Glasso | $0.51 \pm 1.0\text{e-}2$ | $0.02 \pm 2.7\text{e-}4$ | 12 |
| | Dcov | $0.61 \pm 1.3\text{e-}2$ | $0.06 \pm 6.2\text{e-}3$ | 39 |
| | DDC | $0.78 \pm 1.4\text{e-}2$ | $0.10 \pm 1.4\text{e-}2$ | 40 |
| Sync | CCG | $0.56 \pm 5.9\text{e-}2$ | $0.02 \pm 4.2\text{e-}3$ | 2000 |
| | ND | $0.58 \pm 2.1\text{e-}2$ | $0.04 \pm 2.4\text{e-}2$ | 6 |
| | Glasso | $0.51 \pm 6.0\text{e-}4$ | $0.02 \pm 1.7\text{e-}4$ | 82 |
| | Dcov | $0.51 \pm 2.4\text{e-}3$ | $0.02 \pm 2.5\text{e-}4$ | 35 |
|  | **DDC** | **$0.79 \pm 2.7\text{e-}2$** | **$0.11 \pm 1.5\text{e-}2$** | **35** |

**Fig. 2.**
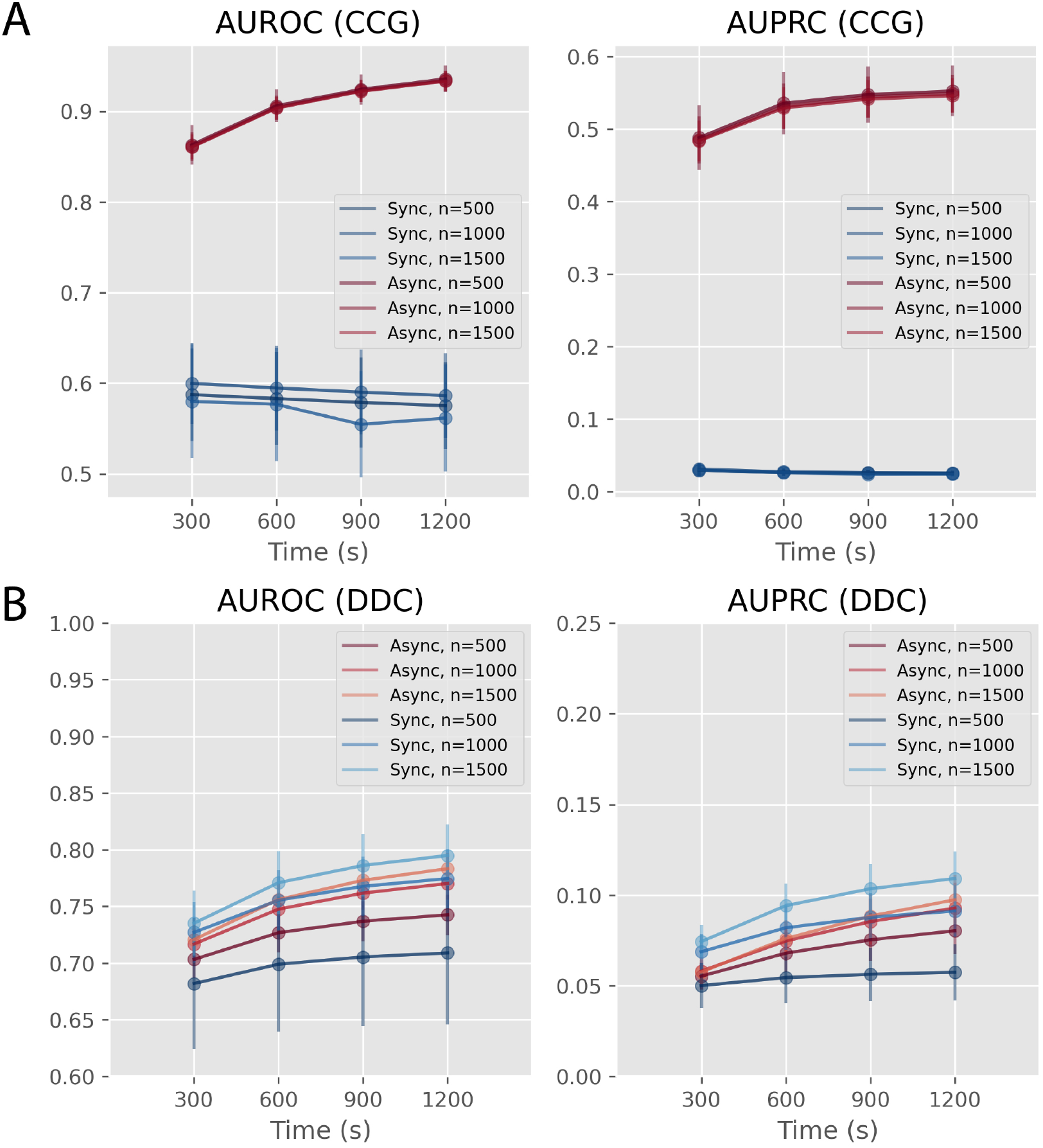
Inference performances of CCG and DDC. The graphs in panels (A) and (B) display average AUROC (left) and AUPRC (right) values (mean *±* standard deviation) across different sampling conditions (number of sampled neurons (n), simulation length used). (A): CCG demonstrated high AUROC and AUPRC values for the asynchronous networks but lower AUROC and AUPRC values in the case of synchronous networks. (B): DDC showed lower average AUROC and AUPRC for the asynchronous networks but showed higher average AUROC and AUPRC values for the synchronous networks compared to CCG.

### Improving the inference performance of Dynamic differential covariance using spike train convolution and fractional differentiation

We further explored the potential of using DDC to infer connections in synchronous networks by increasing the number of data points through computing instantaneous rates. Our findings showed that DDC’s ability to infer connections was enhanced when instantaneous rates were used, with all cases reporting higher average AUPRC values (Fig. 3A, Table 2, S7, S8). Among the four kernels that were used to convolve the spike trains, the Gaussian kernel (*w* =3 ms) consistently showed the highest average AU-ROC and AUPRC scores.

**Table 2.** Performances of modified DDC methods on convolved spike trains. The table shows the performance (AUROC, AUPRC) of modified DDC methods on simulated network activities (1500 sampled neurons and 20 min of simulation). FDDC (*β* = 0.1) with Alpha kernel (*w* = 3 ms) showed the best performance among all combinations (*β, w*) tested in this study. Among the DDC methods utilizing full-integer differentiation, the Gaussian kernel (*w* = 3 ms) emerged as the most effective yet showed lower average AUROC, AUPRC values than the best FDDC method.

| Network act. | Method | Kernel | Filter size<br>($w$, ms) | AUROC | AUPRC | Comp. time (s)<br>(averaged) |
| --- | --- | --- | --- | --- | --- | --- |
| Async | DDC | Alpha | 1 | $0.74 \pm 0.01$ | $0.13 \pm 0.02$ | 290 |
| | | | 3 | $0.79 \pm 0.01$ | $0.11 \pm 0.02$ | 300 |
| | | Gaussian | 1 | $0.75 \pm 0.01$ | $0.13 \pm 0.02$ | 300 |
| | | | 3 | $0.85 \pm 0.02$ | $0.14 \pm 0.02$ | 310 |
| | FDDC ($\beta=0.1$) | Alpha | 1 | $0.79 \pm 0.01$ | $0.17 \pm 0.02$ | 380 |
|  |  |  | 3 | **$0.87 \pm 0.01$** | **$0.31 \pm 0.04$** | **380** |
| | | Gaussian | 1 | $0.80 \pm 0.01$ | $0.18 \pm 0.03$ | 380 |
| | | | 3 | $0.86 \pm 0.01$ | $0.24 \pm 0.03$ | 380 |
| Sync | DDC | Alpha | 1 | $0.75 \pm 0.02$ | $0.14 \pm 0.01$ | 320 |
| | | | 3 | $0.77 \pm 0.02$ | $0.12 \pm 0.01$ | 290 |
| | | Gaussian | 1 | $0.76 \pm 0.02$ | $0.15 \pm 0.01$ | 290 |
| | | | 3 | $0.87 \pm 0.02$ | $0.17 \pm 0.01$ | 290 |
| | FDDC ($\beta=0.1$) | Alpha | 1 | $0.81 \pm 0.02$ | $0.28 \pm 0.02$ | 380 |
|  |  |  | 3 | **$0.92 \pm 0.02$** | **$0.49 \pm 0.03$** | **380** |
| | | Gaussian | 1 | $0.83 \pm 0.02$ | $0.30 \pm 0.02$ | 380 |
| | | | 3 | $0.91 \pm 0.01$ | $0.39 \pm 0.02$ | 380 |

**Fig. 3.**
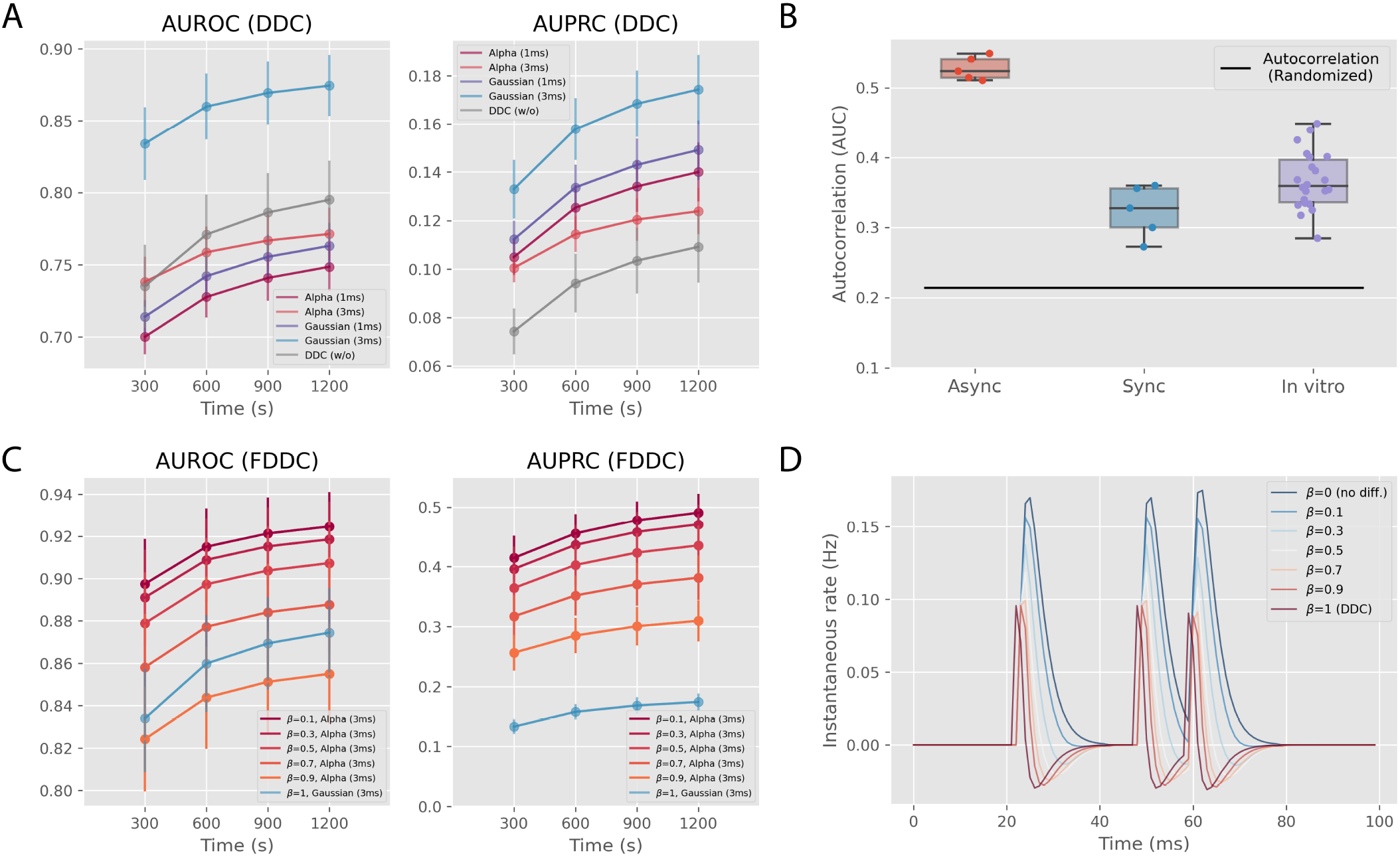
Enhancing inference performance of DDC for synchronous networks (n=1500) (A): AUROC (left) and AUPRC (right) values for the four kernel variants are shown. Gaussian kernel (*w* = 3 ms) consistently showed the best performance. (B): The plot compares the average AUC values of autocorrelation curves for 10 simulated networks (5 asynchronous and 5 synchronous) and 22 in vitro recordings against the average value of random surrogates (standard deviation (*σ*) < 10^*−*5^). For all networks, the average AUC values of autocorrelation curves were larger than those of random surrogates. (C): The performances of FDDC on the synchronous networks for different fractional orders (*β*) are presented (AUROC (left), AUPRC (right)). The results showed that as smaller fractional derivative orders were applied, the average AUROC and AUPRC values increased. The best-performing method from panel A (Gaussian kernel (*w* = 3 ms), colored in cyan) is also plotted for comparison. (D): The plot illustrates how fractional differentiation can preserve the original time series (*β* = 0, no diff.) compared to the full integer differentiation (*β* = 1, the same as DDC). As the *β* increases, the differentiated time series diverges from the original time series.

Our focus then turned to a limitation of the differentiation used in DDC. We noted that the spike trains from simulated network activities and in vitro recordings were non-random processes (e.g., refractory periods). Therefore, we considered fractional differentiation as a means to capture lasting trends in the time series that would be overlooked with full-integer differentiation. Our analysis confirmed that both the simulated and experimental spike trains displayed stronger autocorrelation than the randomized surrogate spike trains (Fig. 3B), indicating the presence of non-random temporal structures in these time series. Compared to DDC using the Gaussian kernel (*w* =3 ms), a consistent improvement in the average AUPRC values was observed for all kernels when fractional differentiation was used (see Table 2, S9, S10). Among these kernels, the Alpha kernel (*w* =3 ms)(see method section “Spike train convolution”) resulted in the best performance (Fig. 3C, Table 2). We also observed that smaller fractional orders (*β*) resulted in enhanced inference performances, owing to better preservation of the original time series with smaller *β* (Fig. 3C, D).

Throughout the study, FDDC (*β* =0.1) combined with Alpha kernel (*w* =3 ms) showed the best performance among all methods that were assessed for the synchronous networks. To test the robustness of this FDDC approach, we probed its performance on the noise-perturbed simulations. We subjected the networks to a broad spectrum of noise levels. At the higher end of the noise range, where the noise level was larger than the leaky current (*≈* 0.2 nA), we observed a recovery of the ground truth connectivity induced by the noise (Table 3). In the lower noise range, we observed consistent performances for both asynchronous and synchronous networks. Regarding average AUROC values, the noise-perturbed asynchronous and synchronous networks showed a decline of approximately 7% and 20%. Similarly, the average AUPRC values of the noise-perturbed asynchronous and synchronous networks displayed a reduction of approximately 32% and 73%, respectively.

**Table 3.**
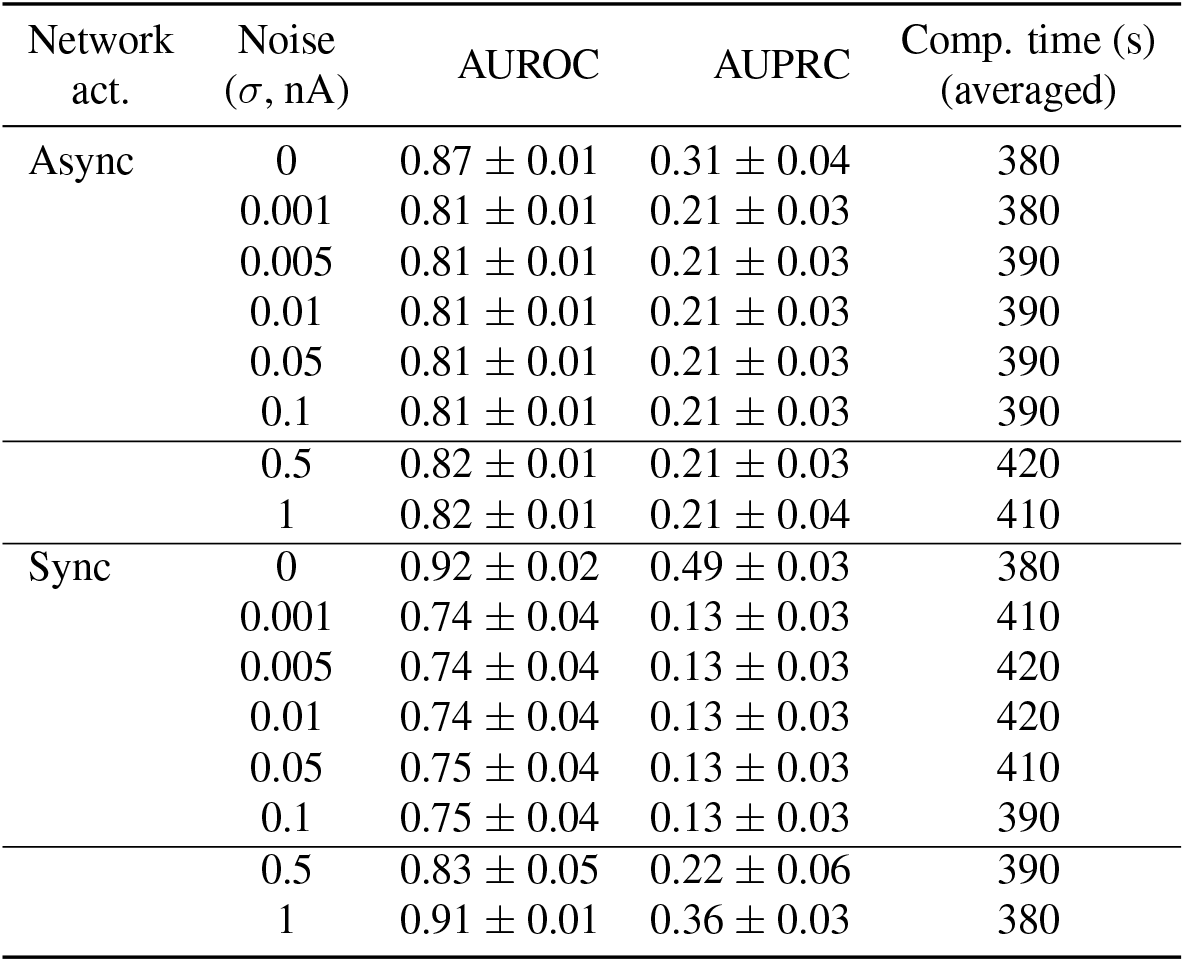
Performances of FDDC (*β* = 0.1, Alpha Kernel (*w* = 3 ms)) on the noise-perturbed simulations. The table shows the performance of FDDC (*β* =0.1) on noiseperturbed simulated networks (1500 sampled neurons and 20 min of simulation). When the noise level was smaller than the leaky current (*≈* 0.2nA), the average AUROC and AUPRC values remained stable. However, in the higher noise range, both values increased, indicating a noise-induced recovery of the ground truth connectivity.

| Network act. | Noise ($\sigma$, nA) | AUROC | AUPRC | Comp. time (s) (averaged) |
| --- | --- | --- | --- | --- |
| Async | 0 | $0.87 \pm 0.01$ | $0.31 \pm 0.04$ | 380 |
| | 0.001 | $0.81 \pm 0.01$ | $0.21 \pm 0.03$ | 380 |
| | 0.005 | $0.81 \pm 0.01$ | $0.21 \pm 0.03$ | 390 |
| | 0.01 | $0.81 \pm 0.01$ | $0.21 \pm 0.03$ | 390 |
| | 0.05 | $0.81 \pm 0.01$ | $0.21 \pm 0.03$ | 390 |
| | 0.1 | $0.81 \pm 0.01$ | $0.21 \pm 0.03$ | 390 |
| | 0.5 | $0.82 \pm 0.01$ | $0.21 \pm 0.03$ | 420 |
| | 1 | $0.82 \pm 0.01$ | $0.21 \pm 0.04$ | 410 |
| Sync | 0 | $0.92 \pm 0.02$ | $0.49 \pm 0.03$ | 380 |
| | 0.001 | $0.74 \pm 0.04$ | $0.13 \pm 0.03$ | 410 |
| | 0.005 | $0.74 \pm 0.04$ | $0.13 \pm 0.03$ | 420 |
| | 0.01 | $0.74 \pm 0.04$ | $0.13 \pm 0.03$ | 420 |
| | 0.05 | $0.75 \pm 0.04$ | $0.13 \pm 0.03$ | 410 |
| | 0.1 | $0.75 \pm 0.04$ | $0.13 \pm 0.03$ | 390 |
| | 0.5 | $0.83 \pm 0.05$ | $0.22 \pm 0.06$ | 390 |
| | 1 | $0.91 \pm 0.01$ | $0.36 \pm 0.03$ | 380 |

### Network connectivity and topology of in vitro neuronal networks differ between FDDC and CCG-derived connectivity graphs

To probe our ability to infer ground truth connectivity using FDDC from synchronous network activity, we inferred connectivity from in vitro hippocampal neuronal cultures using CCG and the best FDDC method. We focused on comparing the resulting graph topology between the two inferred graphs by quantifying the small-world index (SWI) and intersecting edges between them. When comparing the inferred graphs with varying graph densities (e.g., *ρ* = 5, 10, 15, 20 %), CCG-derived graphs generally showed higher SWIs than FDDC-derived graphs (Fig. 4A). For all graph densities, the graphs obtained through CCG consistently exhibited small-worldness with SWI *>* 1, whereas a few of the FDDC-derived graphs displayed SWI *<* 1. Next, we investigated whether there was a relationship between synchronized network firing activities and graph topology by analyzing the correlation between the normalized participation ratio (PR) and SWI. For the graphs derived from FDDC, we found a significant negative linear correlation between the PR and SWI (ordinary least squares, two-sided t-test, p-value < 0.01) across all tested graph densities. This finding shows that the more the connections clustered between the nodes, the more synchronous was the population firing activity of the networks. However, we did not observe any linear correlation for CCG-derived graphs for all tested graph densities (Fig. 4B,C). Finally, we calculated the number of intersecting edges across all graph densities and found that there were always less than 40% of edges that intersected (Fig. 4D-H).

**Fig. 4.**
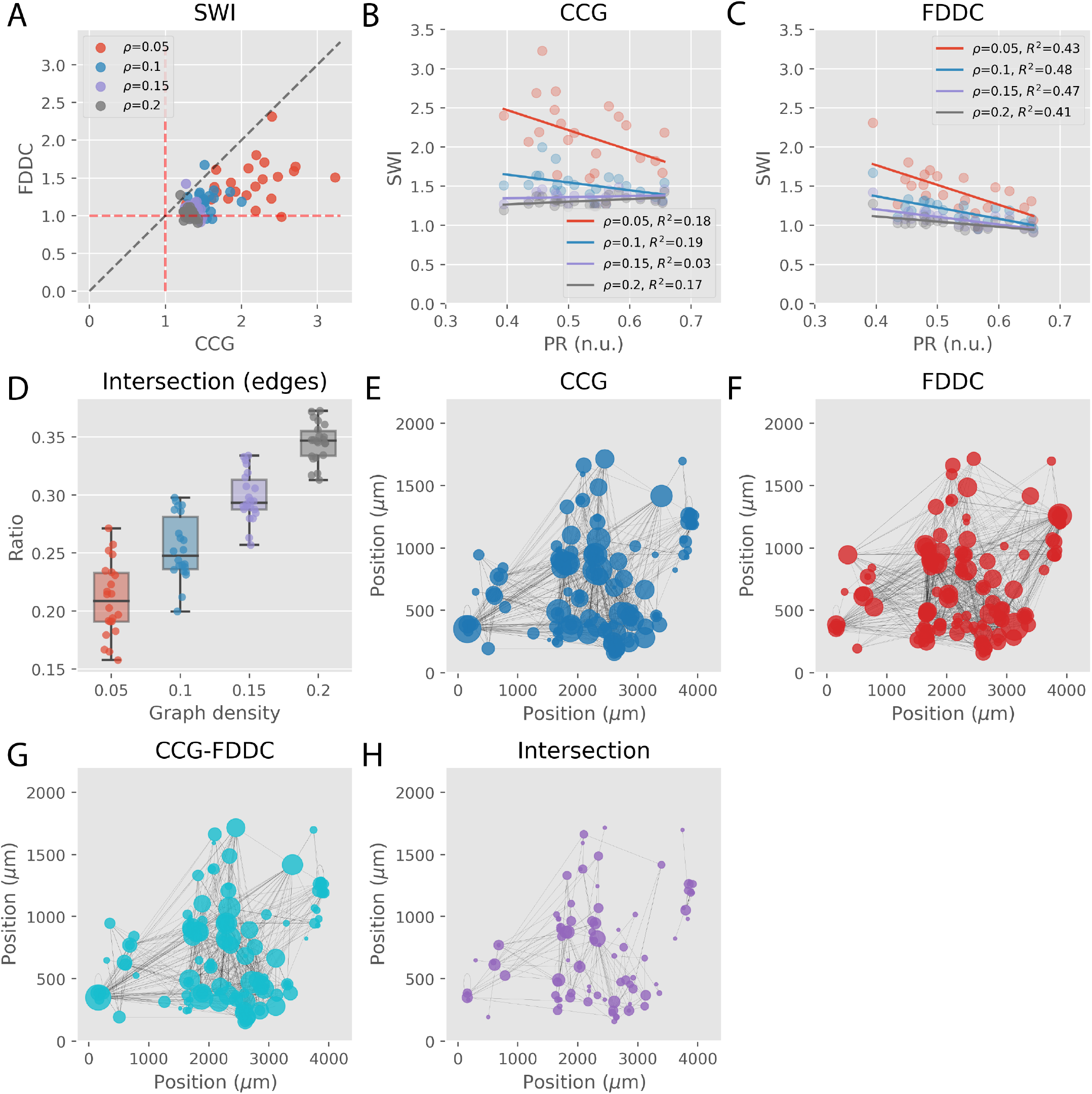
Comparing connectivity graphs of in vitro neuronal networks inferred using CCG and FDDC. (A): The CCG-derived graphs exhibited a small-worldness index (SWI) with values greater than 1 for all the tested graph densities. On the other hand, some of the FDDC-derived graphs had SWI values smaller than 1. In general, the SWI values were higher for CCG-derived graphs compared to FDDC-derived graphs. (B): There was no significant linear correlation between PR and SWI for CCG-derived graphs at any graph density. (C): There was a negative linear correlation between PR and SWI for FDDC-derived graphs across all the graph densities. (D): The ratios of intersecting edges to all existing edges are presented. None of the in vitro neural networks shared more than 40% of edges for all tested graph densities. (E): The panel depicts an example CCG-derived graph, where the size of each node represents the total degree (sum of in-degree and out-degree) of the respective nodes. (F): Panel F depicts the corresponding FDDC-derived graph for the graph shown in (E). (G): Panel G depicts the difference between the CCG-derived graph (panel E) and the FDDC-derived graph (panel F), i.e., only the edges that are exclusive to the CCG-derived graph. (H): Intersection between the CCG-derived graph (panel E) and FDDC-derived graph (panel F) is presented.

## Discussion

In the first part of our study, we compared the performance of selected multivariate inference methods for inferring neuronal connectivity in networks with asynchronous and synchronous activity obtained from LIF simulations. Our findings indicate that when computation time is not a primary concern, the multivariate methods were consistently less effective than CCG for asynchronous networks. Furthermore, we observed subpar performance by ND, Dcov, and Glasso in both asynchronous and synchronous networks. The poor performance of ND may stem from its inability to explicitly consider negative edge weights for the underlying connectivity graph (32). Similar approaches based on graph signal propagation assume undirected and non-negative edges for connectivity graphs (54–57). This assumption may be appropriate for analyzing brain connectivity in data acquired from techniques such as functional magnetic resonance imaging (fMRI), electroencephalography (EEG) and magnetoencephalography (MEG) (58–60). However, the assumption may not accurately reflect inhibitory connections in neuronal cultures. Moreover, Dcov and Glasso, which are both conceptual components of DDC, performed poorly, while DDC emerged as the most effective method on synchronous networks. The findings suggest that accurate recovery of the ground-truth connectivity in synchronous networks requires a combination of directed information obtained through differentiation of the original time series (Dcov) and consideration of other neurons’ impact (Glasso).

We also found that using instantaneous rates to infer connectivity improved the inference performance of DDC. It is worth noting that the idea of convolving spike trains with kernels has been previously proposed by Chen et al. (25), and our study provides quantitative support for this approach. In our experiments, we found that when using full-integer differentiation, the best inference performance was obtained with the Gaussian kernel (*w* = 3 ms). However, when fractional differentiation was employed, the Alpha kernel (*w* = 3 ms) yielded the best performance. This observation suggests that fractional differentiation can preserve useful temporal information that was uniquely captured by an asymmetric kernel. The use of fractional differentiation with a lower fractional order (*β*) improved the inference performance. It should be noted that *β* can be even smaller than the values examined in this study but must be strictly greater than 0; otherwise, we retrieve an identity matrix. In this study, our focus was to demonstrate that FDDC with any *β* outperformed DDC with full-integer differentiation when using instantaneous rates generated with the Alpha kernel (*w* = 3 ms). In practice, we suggest determining the smallest *β* by conducting statistical tests such as the Dickey–Fuller test (61) to ensure stationarity in the differentiated time series. On the other hand, exploring the best-performing FDDC method (*β* = 0.1, Alpha kernel (*w* = 3 ms)) on noise-perturbed networks revealed that the inference using FDDC on synchronous networks was more vulnerable compared to the case of asynchronous networks. This finding suggests FDDC is most effective for networks with robust and consistent synchronized firing patterns.

We compared the inferred connectivity graphs obtained from in vitro neural networks by applying both CCG and FDDC to assess the relevance of FDDC to experimental data. As smallworldness has been previously reported as a characteristic feature of inferred connectivity based on cross-covariance for neuronal cultures (62, 63), we compared the SWI resulting from CCG- and FDDC-derived graphs. In this study, cross-covariance reduces to CCG, as we assumed the original time series were stationary (29), given that FDDC showed improved inference performance over DDC for the tested fractional orders (64). The connectivity graphs obtained using CCG indeed exhibited small-worldness across all tested graph densities. However, some of the connectivity graphs derived using FDDC did not show small-worldness with SWIs smaller than 1, and most of the FDDC-derived graphs showed smaller SWIs compared to CCG-derived graphs. Previous theoretical studies on network dynamics have indicated that networks with clustered connections are more prone to show synchronization (51, 65–67). Our results showed that this relation was robust for FDDC-derived graphs, while no clear correlation was observed for the CCG-derived graphs. We speculate that FDDC-derived graphs generated more conservative connectivity that reflected the synaptic connectivity responsible for the synchronized firing activity.

In conclusion, we propose that the combination of fractional differentiation and DDC presents a promising solution to address the scalability challenges associated with synchronous networks containing thousands of neurons. However, we acknowledge that even the proposed FDDC method would not scale for networks with a much larger number of neurons. The most computationally expensive step in FDDC is the inversion of a large covariance matrix. Therefore, we recommend further exploring methods such as Cholesky decomposition (68) to compute the inverse efficiently or to solve equations directly using proximal gradient methods (69, 70). By implementing these modifications, we believe that derivative inference methods could achieve scalability beyond thousands of neurons.

## Conflict of Interest Statement

The authors declare that the research was conducted in the absence of any commercial or financial relationships that could be construed as a potential conflict of interest.

## Author Contributions

TK, DC, PH, SK contributed to the conception and design of the study. AH, DC, MS, and KB provided scientific supervision to TK. AH and KB provided the necessary resources and materials for the current study. TK performed the experiment and analysis. TK wrote the first draft of the manuscript. All authors contributed to the manuscript revision and approved the submitted version.

## Funding

This work was supported by the European Research Council Advanced Grant 694829 ‘neuroXscales’ and a Swiss Data Science Center project grant (C18-10).

## ACKNOWLEDGEMENTS

We thank Maxwell Biosystems, Zurich, Switzerland, for providing the recording hardware and software support. Dr. Julian Bartram and Xiaohan Xue, all at ETH Zurich, are acknowledged for engaging discussion and feedback.

## Data and code availability

The scripts used in this study are available from the following GitHub repository: https://github.com/arahangua/FDDC.

## Supplementary Material

**Table S1.** LIF simulation parameters. In this study, *Q*_*E*_ and *Q*_*I*_ were fixed as (10 nS, 65 nS) after the parameter search.

| Parameters | Value | Unit |
| --- | --- | --- |
| $C$ (membrane capacitance) | 200 | pF |
| $E_L$ (leaky membrane potential) | -60 | mV |
| $g_L$ (leakage conductance) | 10 | nS |
| $V_T$ (effective threshold) | -50 | mV |
| $V_{\text{reset}}$ (reset potential) | -60 | mV |
| $T_{\text{refrac}}$ (refractory period) | 5 | 5ms |
| $\Delta$ (threshold slope factor) | 2.5 | ms |
| $E_E$ (reversal potential (exc.)) | 0 | mV |
| $E_I$ (reversal potential (inh.)) | -80 | mV |
| $Q_E$ (synaptic strength change (exc.)) | 10 | nS |
| $Q_I$ (synaptic strength change (inh.)) | 65 | nS |
| $\tau_m$ (time constant for white noise) | 20 | ms |

**Table S2.** Parameters for spike-sorting (Kilosort2)

| Parameters | value |
| --- | --- |
| ops.fs | 20000 |
| ops.fshigh | 150 |
| ops.minfr_goodchannels | 0 |
| ops.Th | [10 4] |
| ops.lam | 10 |
| ops.AUCsplit | 0.9 |
| ops.minFR | 1/50 |
| ops.momentum | [20 400] |
| ops.sigmaMask | 30 |
| ops.ThPre | 8 |
| ops.spkTh | -6 |
| ops.reorder | 1 |
| ops.nskip | 25 |
| ops.nfilt_factor | 4 |
| ops.ntbuff | 64 |
| ops.NT | 4*64*1024+ ops.ntbuff |
| ops.whiteningRange | 32 |
| ops.nSkipCov | 25 |
| ops.scaleproc | 200 |
| ops.nPCs | 3 |

**Table S3.** Performance of CCG method. The table shows the performance (AUROC, AUPRC) of CCG method across different inference conditions.

| Method | Network | Neurons | Duration (s) | AUROC |  | AUPRC |  | Comp. time (s) |  |
| --- | --- | --- | --- | --- | --- | --- | --- | --- | --- |
|  |  |  |  | mean | std. | mean | std. | mean | std. |
| CCG | Async | 500 | 300 | 0.86 | 0.022 | 0.49 | 0.044 | 66 | 16 |
|  |  |  | 600 | 0.91 | 0.018 | 0.54 | 0.043 | 110 | 20 |
|  |  |  | 900 | 0.92 | 0.017 | 0.55 | 0.039 | 160 | 36 |
|  |  |  | 1200 | 0.94 | 0.015 | 0.55 | 0.035 | 200 | 42 |
|  |  | 1000 | 300 | 0.86 | 0.015 | 0.48 | 0.032 | 350 | 50 |
|  |  |  | 600 | 0.9 | 0.013 | 0.53 | 0.031 | 630 | 79 |
|  |  |  | 900 | 0.92 | 0.012 | 0.54 | 0.028 | 930 | 140 |
|  |  |  | 1200 | 0.93 | 0.011 | 0.55 | 0.025 | 1200 | 210 |
|  |  | 1500 | 300 | 0.86 | 0.014 | 0.48 | 0.028 | 880 | 46 |
|  |  |  | 600 | 0.9 | 0.013 | 0.53 | 0.028 | 1600 | 150 |
|  |  |  | 900 | 0.92 | 0.012 | 0.54 | 0.026 | 2500 | 420 |
|  |  |  | 1200 | 0.93 | 0.011 | 0.55 | 0.024 | 3300 | 290 |
|  | Sync | 500 | 300 | 0.59 | 0.051 | 0.029 | 0.0082 | 51 | 5.3 |
|  |  |  | 600 | 0.58 | 0.051 | 0.026 | 0.005 | 88 | 12 |
|  |  |  | 900 | 0.58 | 0.05 | 0.025 | 0.0041 | 120 | 17 |
|  |  |  | 1200 | 0.58 | 0.048 | 0.025 | 0.0036 | 150 | 15 |
|  |  | 1000 | 300 | 0.6 | 0.045 | 0.031 | 0.0083 | 250 | 30 |
|  |  |  | 600 | 0.59 | 0.047 | 0.027 | 0.0051 | 470 | 69 |
|  |  |  | 900 | 0.59 | 0.047 | 0.026 | 0.0043 | 680 | 92 |
|  |  |  | 1200 | 0.59 | 0.047 | 0.025 | 0.0038 | 880 | 140 |
|  |  | 1500 | 300 | 0.58 | 0.063 | 0.029 | 0.0093 | 570 | 57 |
|  |  |  | 600 | 0.58 | 0.062 | 0.026 | 0.0058 | 1100 | 200 |
|  |  |  | 900 | 0.55 | 0.059 | 0.024 | 0.0045 | 1600 | 260 |
|  |  |  | 1200 | 0.56 | 0.059 | 0.024 | 0.0042 | 2000 | 250 |

**Table S4.**
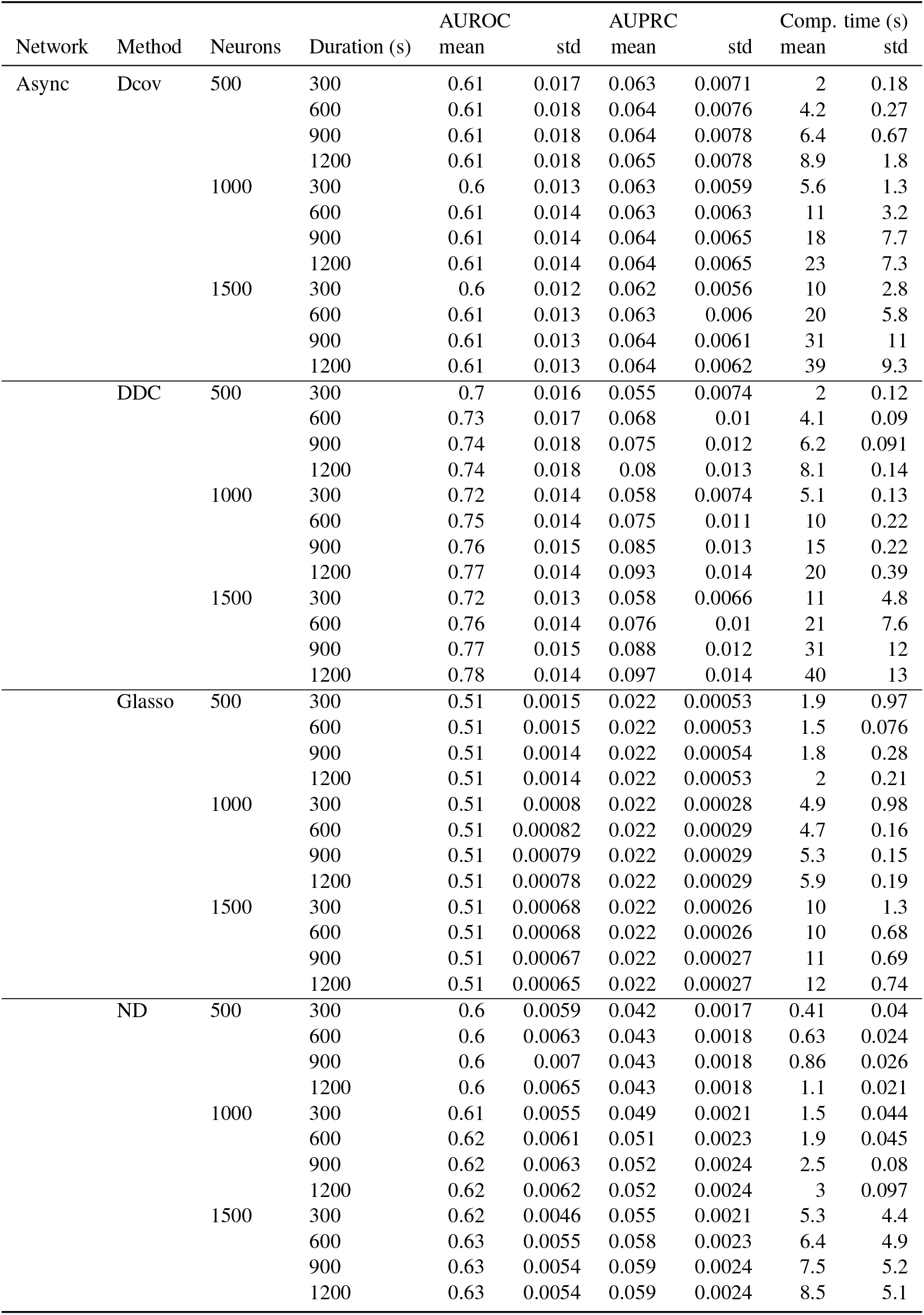
Performance of multivariate methods (asynchronous networks). The table shows the performance (AUROC, AUPRC) of multivariate methods across different inference conditions.

**Table S5.** Performance of multivariate methods (synchronous networks). The table shows the performance (AUROC, AUPRC) of multivariate methods across different inference conditions.

| Network | Method | Neurons | Duration (s) | AUROC |  | AUPRC |  | Comp. time (s) |  |
| --- | --- | --- | --- | --- | --- | --- | --- | --- | --- |
|  |  |  |  | mean | std | mean | std | mean | std |
| Sync | Dcov | 500 | 300 | 0.51 | 0.0056 | 0.021 | 0.00053 | 2 | 0.13 |
|  |  |  | 600 | 0.51 | 0.0054 | 0.021 | 0.00053 | 4.1 | 0.13 |
|  |  |  | 900 | 0.51 | 0.0054 | 0.021 | 0.00053 | 6.1 | 0.23 |
|  |  |  | 1200 | 0.51 | 0.0054 | 0.021 | 0.00052 | 8.2 | 0.27 |
|  |  | 1000 | 300 | 0.51 | 0.0032 | 0.021 | 0.00036 | 5 | 0.075 |
|  |  |  | 600 | 0.51 | 0.003 | 0.021 | 0.00035 | 10 | 0.23 |
|  |  |  | 900 | 0.51 | 0.003 | 0.021 | 0.00035 | 15 | 0.42 |
|  |  |  | 1200 | 0.51 | 0.003 | 0.021 | 0.00035 | 20 | 0.58 |
|  |  | 1500 | 300 | 0.51 | 0.0027 | 0.021 | 0.00026 | 8.8 | 0.12 |
|  |  |  | 600 | 0.51 | 0.0024 | 0.021 | 0.00025 | 18 | 0.32 |
|  |  |  | 900 | 0.51 | 0.0024 | 0.021 | 0.00025 | 26 | 0.56 |
|  |  |  | 1200 | 0.51 | 0.0024 | 0.021 | 0.00025 | 35 | 0.69 |
|  | DDC | 500 | 300 | 0.68 | 0.058 | 0.05 | 0.012 | 2.8 | 1.5 |
|  |  |  | 600 | 0.7 | 0.06 | 0.054 | 0.014 | 5 | 1.9 |
|  |  |  | 900 | 0.71 | 0.061 | 0.056 | 0.015 | 7.1 | 2.3 |
|  |  |  | 1200 | 0.71 | 0.063 | 0.057 | 0.015 | 9.1 | 2.2 |
|  |  | 1000 | 300 | 0.73 | 0.027 | 0.069 | 0.0098 | 6.1 | 2.1 |
|  |  |  | 600 | 0.76 | 0.026 | 0.082 | 0.012 | 13 | 5.4 |
|  |  |  | 900 | 0.77 | 0.026 | 0.088 | 0.013 | 17 | 3.4 |
|  |  |  | 1200 | 0.77 | 0.026 | 0.091 | 0.014 | 23 | 6.1 |
|  |  | 1500 | 300 | 0.74 | 0.029 | 0.074 | 0.0095 | 9 | 0.13 |
|  |  |  | 600 | 0.77 | 0.028 | 0.094 | 0.012 | 18 | 0.3 |
|  |  |  | 900 | 0.79 | 0.027 | 0.1 | 0.014 | 27 | 0.61 |
|  |  |  | 1200 | 0.79 | 0.027 | 0.11 | 0.015 | 35 | 0.48 |
|  | Glasso | 500 | 300 | 0.51 | 0.0026 | 0.021 | 0.0005 | 31 | 54 |
|  |  |  | 600 | 0.51 | 0.0028 | 0.021 | 0.0005 | 28 | 48 |
|  |  |  | 900 | 0.51 | 0.0027 | 0.021 | 0.00049 | 23 | 40 |
|  |  |  | 1200 | 0.51 | 0.0028 | 0.021 | 0.00051 | 23 | 38 |
|  |  | 1000 | 300 | 0.51 | 0.00083 | 0.021 | 0.00031 | 45 | 55 |
|  |  |  | 600 | 0.51 | 0.00087 | 0.021 | 0.00031 | 35 | 42 |
|  |  |  | 900 | 0.51 | 0.00086 | 0.021 | 0.00032 | 53 | 71 |
|  |  |  | 1200 | 0.51 | 0.00087 | 0.021 | 0.00032 | 53 | 73 |
|  |  | 1500 | 300 | 0.51 | 0.00056 | 0.021 | 0.00017 | 70 | 71 |
|  |  |  | 600 | 0.51 | 0.00055 | 0.021 | 0.00017 | 80 | 91 |
|  |  |  | 900 | 0.51 | 0.00055 | 0.021 | 0.00017 | 78 | 86 |
|  |  |  | 1200 | 0.51 | 0.00056 | 0.021 | 0.00017 | 82 | 93 |
|  | ND | 500 | 300 | 0.55 | 0.021 | 0.028 | 0.0033 | 1.2 | 1.6 |
|  |  |  | 600 | 0.55 | 0.021 | 0.028 | 0.0034 | 1.4 | 1.7 |
|  |  |  | 900 | 0.55 | 0.021 | 0.028 | 0.0034 | 1.6 | 1.5 |
|  |  |  | 1200 | 0.55 | 0.021 | 0.028 | 0.0034 | 1.8 | 1.6 |
|  |  | 1000 | 300 | 0.57 | 0.017 | 0.033 | 0.0037 | 2.7 | 2.4 |
|  |  |  | 600 | 0.57 | 0.017 | 0.034 | 0.0037 | 3.5 | 3 |
|  |  |  | 900 | 0.57 | 0.017 | 0.034 | 0.0038 | 4 | 3 |
|  |  |  | 1200 | 0.57 | 0.016 | 0.034 | 0.0037 | 4 | 1.9 |
|  |  | 1500 | 300 | 0.57 | 0.021 | 0.035 | 0.0049 | 3.3 | 0.083 |
|  |  |  | 600 | 0.58 | 0.021 | 0.036 | 0.0052 | 4.2 | 0.046 |
|  |  |  | 900 | 0.58 | 0.021 | 0.036 | 0.0053 | 5.3 | 0.23 |
|  |  |  | 1200 | 0.58 | 0.021 | 0.037 | 0.0053 | 6.2 | 0.17 |

**Table S6.** Performance of undirected multivariate methods. The table shows the performance (AUROC, AUPRC) of the undirected multivariate methods (Glasso, ND) evaluated using the undirected ground truth connections.

| Network | Method | Neurons | Duration (s) | AUROC |  | AUPRC |  | Comp. time (s) |  |
| --- | --- | --- | --- | --- | --- | --- | --- | --- | --- |
|  |  |  |  | mean | std | mean | std | mean | std |
| Async | Glasso | 500 | 300 | 0.51 | 0.0015 | 0.043 | 0.00089 | 1.9 | 0.97 |
|  |  |  | 600 | 0.51 | 0.0015 | 0.043 | 0.00088 | 1.5 | 0.076 |
|  |  |  | 900 | 0.51 | 0.0014 | 0.043 | 0.00089 | 1.8 | 0.28 |
|  |  |  | 1200 | 0.51 | 0.0014 | 0.043 | 0.00088 | 2 | 0.21 |
|  |  | 1000 | 300 | 0.51 | 0.00081 | 0.043 | 0.00054 | 4.9 | 0.98 |
|  |  |  | 600 | 0.51 | 0.00083 | 0.043 | 0.00055 | 4.7 | 0.16 |
|  |  |  | 900 | 0.51 | 0.00079 | 0.043 | 0.00055 | 5.3 | 0.15 |
|  |  |  | 1200 | 0.51 | 0.00079 | 0.043 | 0.00054 | 5.9 | 0.19 |
|  |  | 1500 | 300 | 0.51 | 0.00066 | 0.044 | 0.00044 | 10 | 1.3 |
|  |  |  | 600 | 0.51 | 0.00066 | 0.044 | 0.00045 | 10 | 0.68 |
|  |  |  | 900 | 0.51 | 0.00065 | 0.044 | 0.00046 | 11 | 0.69 |
|  |  |  | 1200 | 0.51 | 0.00064 | 0.044 | 0.00046 | 12 | 0.74 |
|  | ND | 500 | 300 | 0.6 | 0.006 | 0.082 | 0.0032 | 0.41 | 0.04 |
|  |  |  | 600 | 0.6 | 0.0064 | 0.084 | 0.0034 | 0.63 | 0.024 |
|  |  |  | 900 | 0.61 | 0.007 | 0.084 | 0.0034 | 0.86 | 0.026 |
|  |  |  | 1200 | 0.61 | 0.0066 | 0.085 | 0.0035 | 1.1 | 0.021 |
|  |  | 1000 | 300 | 0.61 | 0.0057 | 0.097 | 0.0042 | 1.5 | 0.044 |
|  |  |  | 600 | 0.62 | 0.0063 | 0.1 | 0.0045 | 1.9 | 0.045 |
|  |  |  | 900 | 0.62 | 0.0065 | 0.1 | 0.0046 | 2.5 | 0.08 |
|  |  |  | 1200 | 0.62 | 0.0063 | 0.1 | 0.0047 | 3 | 0.097 |
|  |  | 1500 | 300 | 0.62 | 0.0047 | 0.11 | 0.0041 | 5.3 | 4.4 |
|  |  |  | 600 | 0.63 | 0.0057 | 0.11 | 0.0047 | 6.4 | 4.9 |
|  |  |  | 900 | 0.63 | 0.0056 | 0.11 | 0.0047 | 7.5 | 5.2 |
|  |  |  | 1200 | 0.63 | 0.0056 | 0.12 | 0.0048 | 8.5 | 5.1 |
| Sync | Glasso | 500 | 300 | 0.51 | 0.0026 | 0.042 | 0.00089 | 31 | 54 |
|  |  |  | 600 | 0.51 | 0.0028 | 0.042 | 0.00093 | 28 | 48 |
|  |  |  | 900 | 0.51 | 0.0028 | 0.042 | 0.00092 | 23 | 40 |
|  |  |  | 1200 | 0.51 | 0.0028 | 0.042 | 0.00094 | 23 | 38 |
|  |  | 1000 | 300 | 0.51 | 0.00083 | 0.042 | 0.00058 | 45 | 55 |
|  |  |  | 600 | 0.51 | 0.00086 | 0.042 | 0.00058 | 35 | 42 |
|  |  |  | 900 | 0.51 | 0.00085 | 0.042 | 0.00058 | 53 | 71 |
|  |  |  | 1200 | 0.51 | 0.00086 | 0.042 | 0.00058 | 53 | 73 |
|  |  | 1500 | 300 | 0.51 | 0.00059 | 0.042 | 0.00033 | 70 | 71 |
|  |  |  | 600 | 0.51 | 0.00056 | 0.042 | 0.00032 | 80 | 91 |
|  |  |  | 900 | 0.51 | 0.00057 | 0.042 | 0.00033 | 78 | 86 |
|  |  |  | 1200 | 0.51 | 0.00058 | 0.042 | 0.00032 | 82 | 93 |
|  | ND | 500 | 300 | 0.55 | 0.021 | 0.055 | 0.0064 | 1.2 | 1.6 |
|  |  |  | 600 | 0.55 | 0.021 | 0.056 | 0.0065 | 1.4 | 1.7 |
|  |  |  | 900 | 0.55 | 0.021 | 0.056 | 0.0065 | 1.6 | 1.5 |
|  |  |  | 1200 | 0.55 | 0.021 | 0.056 | 0.0065 | 1.8 | 1.6 |
|  |  | 1000 | 300 | 0.57 | 0.017 | 0.065 | 0.0071 | 2.7 | 2.4 |
|  |  |  | 600 | 0.57 | 0.017 | 0.066 | 0.0072 | 3.5 | 3 |
|  |  |  | 900 | 0.57 | 0.017 | 0.067 | 0.0073 | 4 | 3 |
|  |  |  | 1200 | 0.57 | 0.017 | 0.067 | 0.0073 | 4 | 1.9 |
|  |  | 1500 | 300 | 0.58 | 0.021 | 0.069 | 0.0096 | 3.3 | 0.083 |
|  |  |  | 600 | 0.58 | 0.022 | 0.071 | 0.01 | 4.2 | 0.046 |
|  |  |  | 900 | 0.58 | 0.022 | 0.072 | 0.01 | 5.3 | 0.23 |
|  |  |  | 1200 | 0.58 | 0.022 | 0.072 | 0.01 | 6.2 | 0.17 |

**Table S7.** Performance of DDC on convolved spike trains (asynchronous networks). The performance (AUROC, AUPRC) of DDC on convolved spike trains is presented in the table for four different kernels.

| Network | Method | Neurons | Duration (s) | AUROC |  | AUPRC |  | Comp. time (s) |  |
| --- | --- | --- | --- | --- | --- | --- | --- | --- | --- |
|  |  |  |  | mean | std | mean | std | mean | std |
| Async | DDC (Alpha, 1 ms) | 500 | 300 | 0.68 | 0.016 | 0.065 | 0.011 | 19 | 0.49 |
|  |  |  | 600 | 0.71 | 0.018 | 0.092 | 0.018 | 39 | 1.4 |
|  |  |  | 900 | 0.73 | 0.018 | 0.11 | 0.023 | 59 | 1.2 |
|  |  |  | 1200 | 0.74 | 0.019 | 0.12 | 0.026 | 79 | 0.91 |
|  |  | 1000 | 300 | 0.68 | 0.013 | 0.065 | 0.0097 | 44 | 0.62 |
|  |  |  | 600 | 0.71 | 0.014 | 0.092 | 0.016 | 88 | 0.74 |
|  |  |  | 900 | 0.73 | 0.015 | 0.11 | 0.02 | 130 | 0.93 |
|  |  |  | 1200 | 0.74 | 0.015 | 0.13 | 0.023 | 180 | 3.1 |
|  |  | 1500 | 300 | 0.68 | 0.011 | 0.064 | 0.0087 | 72 | 3.6 |
|  |  |  | 600 | 0.71 | 0.012 | 0.092 | 0.015 | 150 | 3.7 |
|  |  |  | 900 | 0.73 | 0.012 | 0.11 | 0.019 | 220 | 4.4 |
|  |  |  | 1200 | 0.74 | 0.013 | 0.13 | 0.022 | 290 | 5.3 |
|  | DDC (Alpha, 3 ms) | 500 | 300 | 0.72 | 0.017 | 0.066 | 0.011 | 20 | 0.76 |
|  |  |  | 600 | 0.75 | 0.018 | 0.079 | 0.014 | 40 | 1.2 |
|  |  |  | 900 | 0.76 | 0.018 | 0.087 | 0.016 | 61 | 1.1 |
|  |  |  | 1200 | 0.76 | 0.017 | 0.092 | 0.018 | 82 | 1.2 |
|  |  | 1000 | 300 | 0.73 | 0.013 | 0.069 | 0.01 | 45 | 1.2 |
|  |  |  | 600 | 0.76 | 0.014 | 0.087 | 0.014 | 90 | 2.5 |
|  |  |  | 900 | 0.77 | 0.014 | 0.097 | 0.016 | 130 | 3.7 |
|  |  |  | 1200 | 0.78 | 0.014 | 0.1 | 0.018 | 180 | 5 |
|  |  | 1500 | 300 | 0.73 | 0.012 | 0.068 | 0.0088 | 76 | 3 |
|  |  |  | 600 | 0.77 | 0.013 | 0.088 | 0.012 | 150 | 4 |
|  |  |  | 900 | 0.78 | 0.013 | 0.1 | 0.014 | 230 | 4.7 |
|  |  |  | 1200 | 0.79 | 0.013 | 0.11 | 0.016 | 300 | 7.5 |
|  | DDC (Gaussian, 1 ms) | 500 | 300 | 0.68 | 0.016 | 0.067 | 0.012 | 20 | 0.67 |
|  |  |  | 600 | 0.72 | 0.018 | 0.095 | 0.019 | 40 | 1.2 |
|  |  |  | 900 | 0.73 | 0.018 | 0.11 | 0.023 | 59 | 2 |
|  |  |  | 1200 | 0.74 | 0.018 | 0.13 | 0.026 | 80 | 1.4 |
|  |  | 1000 | 300 | 0.69 | 0.014 | 0.068 | 0.01 | 44 | 0.55 |
|  |  |  | 600 | 0.72 | 0.014 | 0.096 | 0.017 | 89 | 1.4 |
|  |  |  | 900 | 0.74 | 0.015 | 0.12 | 0.021 | 130 | 2 |
|  |  |  | 1200 | 0.75 | 0.015 | 0.13 | 0.024 | 180 | 1.8 |
|  |  | 1500 | 300 | 0.69 | 0.011 | 0.067 | 0.0092 | 74 | 2.5 |
|  |  |  | 600 | 0.72 | 0.012 | 0.096 | 0.016 | 150 | 3.5 |
|  |  |  | 900 | 0.74 | 0.012 | 0.12 | 0.02 | 220 | 5.6 |
|  |  |  | 1200 | 0.75 | 0.013 | 0.13 | 0.023 | 300 | 6.7 |
|  | DDC (Gaussian, 3 ms) | 500 | 300 | 0.76 | 0.02 | 0.073 | 0.012 | 21 | 0.41 |
|  |  |  | 600 | 0.78 | 0.021 | 0.086 | 0.015 | 41 | 0.66 |
|  |  |  | 900 | 0.78 | 0.021 | 0.093 | 0.016 | 62 | 1.2 |
|  |  |  | 1200 | 0.79 | 0.021 | 0.097 | 0.017 | 83 | 1.7 |
|  |  | 1000 | 300 | 0.79 | 0.016 | 0.084 | 0.013 | 46 | 1.3 |
|  |  |  | 600 | 0.81 | 0.017 | 0.1 | 0.017 | 92 | 1.1 |
|  |  |  | 900 | 0.82 | 0.016 | 0.11 | 0.019 | 140 | 1.9 |
|  |  |  | 1200 | 0.83 | 0.016 | 0.12 | 0.02 | 180 | 2.2 |
|  |  | 1500 | 300 | 0.8 | 0.015 | 0.088 | 0.012 | 77 | 2.7 |
|  |  |  | 600 | 0.83 | 0.015 | 0.11 | 0.016 | 150 | 2.9 |
|  |  |  | 900 | 0.84 | 0.015 | 0.13 | 0.018 | 230 | 4.1 |
|  |  |  | 1200 | 0.85 | 0.015 | 0.14 | 0.019 | 310 | 6.6 |

**Table S8.** Performance of DDC on convolved spike trains (synchronous networks). The performance (AUROC, AUPRC) of DDC on convolved spike trains is presented in the table for four different kernels.

| Network | Method | Neurons | Duration (s) | AUROC |  | AUPRC |  | Comp. time (s) |  |
| --- | --- | --- | --- | --- | --- | --- | --- | --- | --- |
|  |  |  |  | mean | std | mean | std | mean | std |
| Sync | DDC (Alpha, 1 ms) | 500 | 300 | 0.66 | 0.045 | 0.074 | 0.018 | 20 | 3 |
|  |  |  | 600 | 0.68 | 0.049 | 0.081 | 0.021 | 41 | 6.4 |
|  |  |  | 900 | 0.68 | 0.05 | 0.083 | 0.021 | 62 | 9 |
|  |  |  | 1200 | 0.69 | 0.052 | 0.085 | 0.022 | 81 | 8.7 |
|  |  | 1000 | 300 | 0.69 | 0.012 | 0.099 | 0.0089 | 43 | 1.6 |
|  |  |  | 600 | 0.72 | 0.014 | 0.11 | 0.011 | 84 | 2.6 |
|  |  |  | 900 | 0.73 | 0.016 | 0.12 | 0.013 | 130 | 4.9 |
|  |  |  | 1200 | 0.73 | 0.017 | 0.12 | 0.014 | 170 | 5.8 |
|  |  | 1500 | 300 | 0.7 | 0.012 | 0.1 | 0.0074 | 71 | 2.6 |
|  |  |  | 600 | 0.73 | 0.014 | 0.13 | 0.0092 | 150 | 13 |
|  |  |  | 900 | 0.74 | 0.016 | 0.13 | 0.011 | 230 | 35 |
|  |  |  | 1200 | 0.75 | 0.017 | 0.14 | 0.012 | 320 | 62 |
|  | DDC (Alpha, 3 ms) | 500 | 300 | 0.68 | 0.052 | 0.062 | 0.013 | 19 | 0.68 |
|  |  |  | 600 | 0.69 | 0.053 | 0.065 | 0.014 | 37 | 1.4 |
|  |  |  | 900 | 0.7 | 0.054 | 0.066 | 0.015 | 56 | 2.1 |
|  |  |  | 1200 | 0.7 | 0.055 | 0.067 | 0.015 | 75 | 3 |
|  |  | 1000 | 300 | 0.73 | 0.017 | 0.09 | 0.0065 | 43 | 0.77 |
|  |  |  | 600 | 0.74 | 0.017 | 0.099 | 0.0075 | 86 | 1.8 |
|  |  |  | 900 | 0.75 | 0.017 | 0.1 | 0.0083 | 130 | 3.2 |
|  |  |  | 1200 | 0.75 | 0.018 | 0.1 | 0.0088 | 170 | 3.2 |
|  |  | 1500 | 300 | 0.74 | 0.017 | 0.1 | 0.006 | 72 | 3.2 |
|  |  |  | 600 | 0.76 | 0.018 | 0.11 | 0.0074 | 140 | 7.6 |
|  |  |  | 900 | 0.77 | 0.018 | 0.12 | 0.0087 | 220 | 9.8 |
|  |  |  | 1200 | 0.77 | 0.018 | 0.12 | 0.0095 | 290 | 13 |
|  | DDC (Gaussian, 1 ms) | 500 | 300 | 0.67 | 0.048 | 0.078 | 0.019 | 19 | 0.39 |
|  |  |  | 600 | 0.69 | 0.051 | 0.085 | 0.022 | 39 | 0.41 |
|  |  |  | 900 | 0.7 | 0.053 | 0.088 | 0.023 | 58 | 0.74 |
|  |  |  | 1200 | 0.7 | 0.055 | 0.09 | 0.023 | 77 | 0.82 |
|  |  | 1000 | 300 | 0.71 | 0.012 | 0.11 | 0.0091 | 42 | 0.88 |
|  |  |  | 600 | 0.73 | 0.014 | 0.12 | 0.012 | 85 | 1.4 |
|  |  |  | 900 | 0.74 | 0.016 | 0.13 | 0.013 | 130 | 2.4 |
|  |  |  | 1200 | 0.75 | 0.016 | 0.13 | 0.014 | 170 | 3.7 |
|  |  | 1500 | 300 | 0.71 | 0.012 | 0.11 | 0.0077 | 77 | 15 |
|  |  |  | 600 | 0.74 | 0.014 | 0.13 | 0.0093 | 160 | 33 |
|  |  |  | 900 | 0.76 | 0.015 | 0.14 | 0.011 | 240 | 55 |
|  |  |  | 1200 | 0.76 | 0.016 | 0.15 | 0.012 | 290 | 4.1 |
|  | DDC (Gaussian, 3 ms) | 500 | 300 | 0.75 | 0.069 | 0.077 | 0.019 | 20 | 0.61 |
|  |  |  | 600 | 0.76 | 0.071 | 0.083 | 0.02 | 39 | 1.1 |
|  |  |  | 900 | 0.77 | 0.072 | 0.085 | 0.021 | 59 | 1.8 |
|  |  |  | 1200 | 0.77 | 0.073 | 0.086 | 0.022 | 78 | 2.5 |
|  |  | 1000 | 300 | 0.82 | 0.023 | 0.12 | 0.011 | 48 | 9 |
|  |  |  | 600 | 0.84 | 0.021 | 0.13 | 0.012 | 98 | 20 |
|  |  |  | 900 | 0.85 | 0.02 | 0.14 | 0.012 | 150 | 27 |
|  |  |  | 1200 | 0.85 | 0.02 | 0.14 | 0.013 | 190 | 34 |
|  |  | 1500 | 300 | 0.83 | 0.025 | 0.13 | 0.012 | 72 | 2.4 |
|  |  |  | 600 | 0.86 | 0.023 | 0.16 | 0.013 | 150 | 7.1 |
|  |  |  | 900 | 0.87 | 0.022 | 0.17 | 0.014 | 220 | 13 |
|  |  |  | 1200 | 0.87 | 0.021 | 0.17 | 0.014 | 290 | 15 |

**Table S9.**
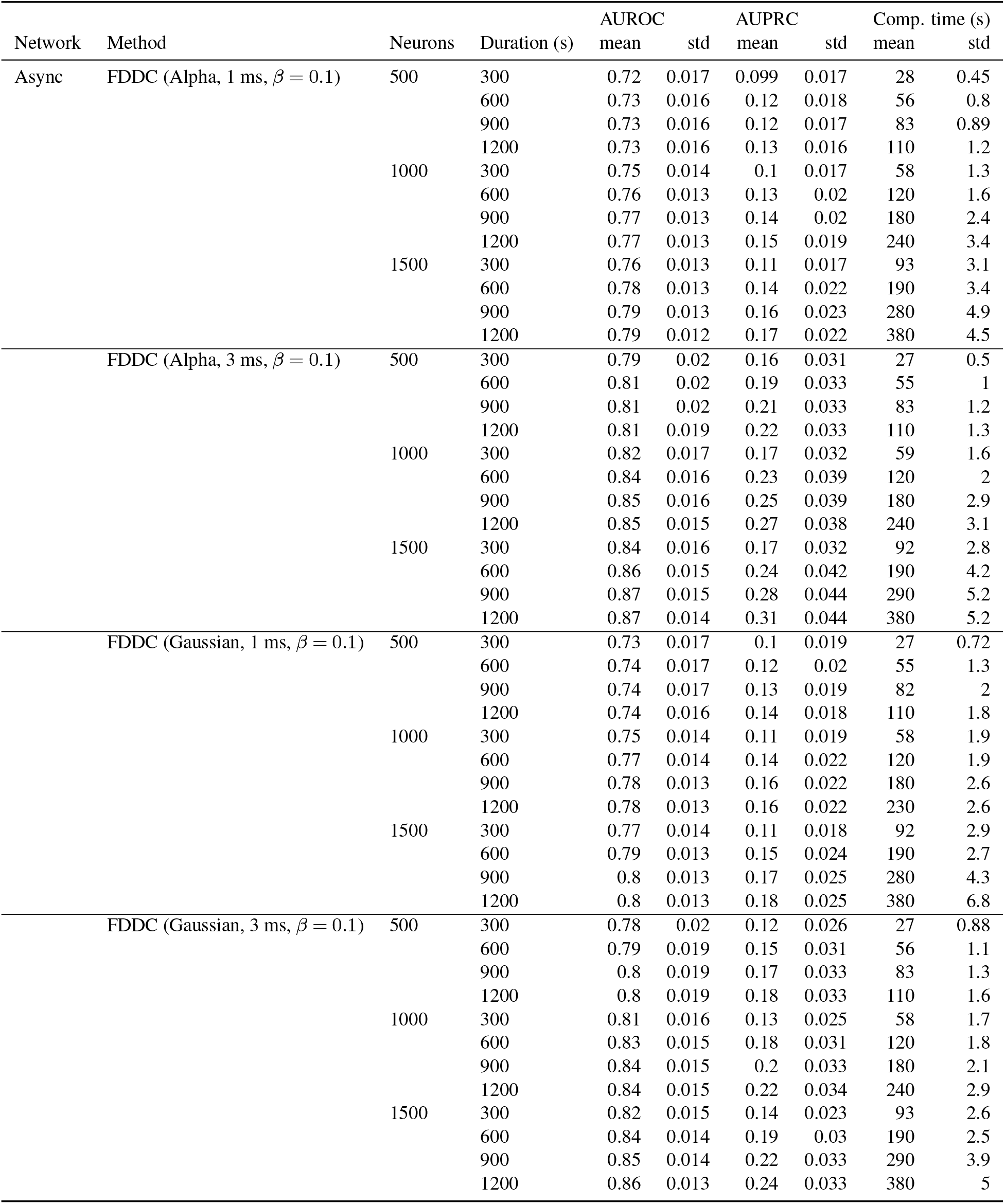
Performance of FDDC on convolved spike trains (asynchronous networks). The performance (AUROC, AUPRC) of FDDC (*β* = 0.1) on convolved spike trains is presented in the table for four different kernels.

**Table S10.** Performance of FDDC on convolved spike trains (synchronous networks). The performance (AUROC, AUPRC) of FDDC (*β* = 0.1) on convolved spike trains is presented in the table for four different kernels.

| Network | Method | Neurons | Duration (s) | AUROC |  | AUPRC |  | Comp. time (s) |  |
| --- | --- | --- | --- | --- | --- | --- | --- | --- | --- |
|  |  |  |  | mean | std | mean | std | mean | std |
| Sync | FDDC (Alpha, 1 ms, $\beta = 0.1$) | 500 | 300 | 0.73 | 0.065 | 0.14 | 0.043 | 27 | 0.47 |
|  |  |  | 600 | 0.74 | 0.066 | 0.15 | 0.045 | 55 | 0.81 |
|  |  |  | 900 | 0.74 | 0.067 | 0.15 | 0.046 | 82 | 1.3 |
|  |  |  | 1200 | 0.74 | 0.067 | 0.15 | 0.046 | 110 | 2.1 |
|  |  | 1000 | 300 | 0.78 | 0.018 | 0.21 | 0.017 | 58 | 1.5 |
|  |  |  | 600 | 0.79 | 0.018 | 0.23 | 0.017 | 120 | 1.6 |
|  |  |  | 900 | 0.8 | 0.018 | 0.24 | 0.016 | 180 | 2.1 |
|  |  |  | 1200 | 0.8 | 0.017 | 0.24 | 0.015 | 240 | 2.1 |
|  |  | 1500 | 300 | 0.79 | 0.019 | 0.23 | 0.018 | 92 | 2.2 |
|  |  |  | 600 | 0.8 | 0.018 | 0.26 | 0.02 | 190 | 1.9 |
|  |  |  | 900 | 0.81 | 0.018 | 0.27 | 0.019 | 280 | 3.2 |
|  |  |  | 1200 | 0.81 | 0.018 | 0.28 | 0.016 | 380 | 4.6 |
| | FDDC (Alpha, 3 ms, $\beta = 0.1$) | 500 | 300 | 0.81 | 0.086 | 0.24 | 0.078 | 27 | 0.37 |
|  |  |  | 600 | 0.82 | 0.088 | 0.25 | 0.081 | 55 | 0.85 |
|  |  |  | 900 | 0.83 | 0.088 | 0.26 | 0.083 | 82 | 0.75 |
|  |  |  | 1200 | 0.83 | 0.089 | 0.26 | 0.084 | 110 | 0.7 |
|  |  | 1000 | 300 | 0.88 | 0.02 | 0.37 | 0.032 | 58 | 1.5 |
|  |  |  | 600 | 0.9 | 0.018 | 0.4 | 0.03 | 120 | 1.6 |
|  |  |  | 900 | 0.9 | 0.017 | 0.42 | 0.03 | 180 | 2.5 |
|  |  |  | 1200 | 0.91 | 0.017 | 0.42 | 0.029 | 240 | 3.5 |
|  |  | 1500 | 300 | 0.9 | 0.021 | 0.41 | 0.037 | 93 | 2.8 |
|  |  |  | 600 | 0.92 | 0.018 | 0.46 | 0.033 | 190 | 2.9 |
|  |  |  | 900 | 0.92 | 0.017 | 0.48 | 0.032 | 280 | 3.7 |
|  |  |  | 1200 | 0.92 | 0.016 | 0.49 | 0.031 | 380 | 4.9 |
| | FDDC (Gaussian, 1 ms, $\beta = 0.1$) | 500 | 300 | 0.74 | 0.068 | 0.16 | 0.047 | 27 | 0.71 |
|  |  |  | 600 | 0.75 | 0.069 | 0.16 | 0.05 | 55 | 1.4 |
|  |  |  | 900 | 0.75 | 0.07 | 0.17 | 0.051 | 82 | 1.9 |
|  |  |  | 1200 | 0.76 | 0.07 | 0.17 | 0.051 | 110 | 2.6 |
|  |  | 1000 | 300 | 0.79 | 0.018 | 0.23 | 0.018 | 58 | 1.9 |
|  |  |  | 600 | 0.8 | 0.018 | 0.25 | 0.018 | 120 | 2.5 |
|  |  |  | 900 | 0.81 | 0.017 | 0.26 | 0.018 | 180 | 3.6 |
|  |  |  | 1200 | 0.81 | 0.017 | 0.26 | 0.016 | 240 | 4.4 |
|  |  | 1500 | 300 | 0.8 | 0.019 | 0.25 | 0.02 | 93 | 2.7 |
|  |  |  | 600 | 0.82 | 0.018 | 0.28 | 0.021 | 190 | 3.5 |
|  |  |  | 900 | 0.82 | 0.018 | 0.29 | 0.02 | 280 | 4 |
|  |  |  | 1200 | 0.83 | 0.018 | 0.3 | 0.018 | 380 | 3.7 |
| | FDDC (Gaussian, 3 ms, $\beta = 0.1$) | 500 | 300 | 0.79 | 0.081 | 0.2 | 0.062 | 27 | 0.53 |
|  |  |  | 600 | 0.8 | 0.082 | 0.21 | 0.066 | 55 | 0.75 |
|  |  |  | 900 | 0.81 | 0.083 | 0.22 | 0.067 | 83 | 0.65 |
|  |  |  | 1200 | 0.81 | 0.084 | 0.22 | 0.068 | 110 | 0.8 |
|  |  | 1000 | 300 | 0.86 | 0.018 | 0.29 | 0.022 | 58 | 1.4 |
|  |  |  | 600 | 0.88 | 0.016 | 0.32 | 0.019 | 120 | 1.3 |
|  |  |  | 900 | 0.89 | 0.015 | 0.34 | 0.019 | 180 | 2.1 |
|  |  |  | 1200 | 0.89 | 0.014 | 0.34 | 0.018 | 240 | 2.7 |
|  |  | 1500 | 300 | 0.88 | 0.02 | 0.32 | 0.025 | 94 | 2.5 |
|  |  |  | 600 | 0.9 | 0.017 | 0.36 | 0.021 | 190 | 3.7 |
|  |  |  | 900 | 0.91 | 0.015 | 0.38 | 0.02 | 290 | 4.6 |
|  |  |  | 1200 | 0.91 | 0.014 | 0.39 | 0.02 | 380 | 6.6 |

**Fig. S1.**
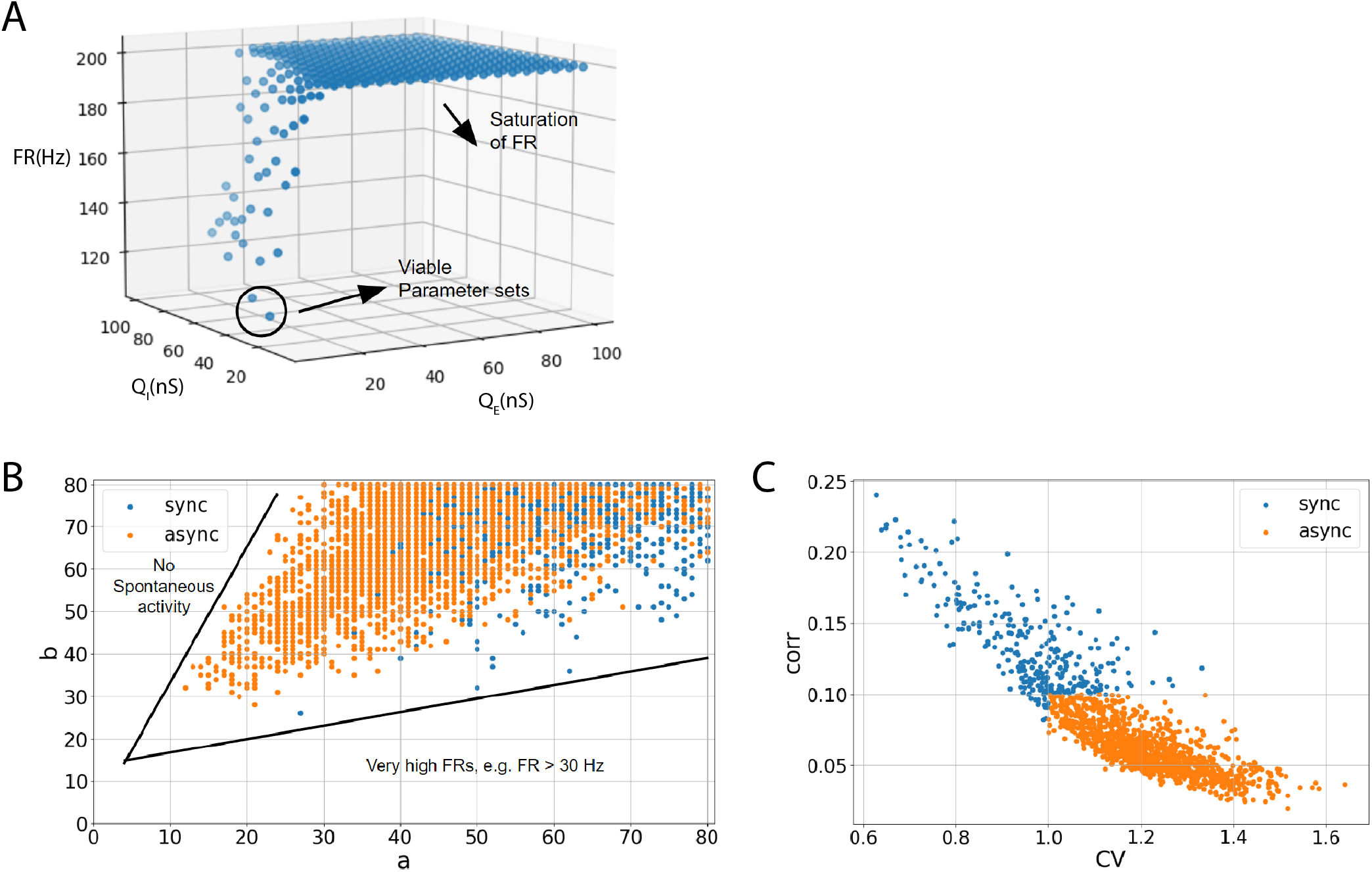
Parameter search of LIF network simulations. (A): While fixing the adaptation parameters (a,b) = (1 *µS*,5*nA*) for excitatory neurons to implement ‘weakly adapting cells’ (24), the amounts for postsynaptic excitatory/inhibitory conductance change (*QE,I*) were searched in the range of [0 100] with a step size of 5. Each parameter set was simulated for 5 seconds, and the corresponding average population firing rates are shown. (B): Under the fixed parameter choice, (*QE, QI*) = (10, 65) *nS*, (a,b) parameter sets were searched in the range [0 80] with a step size of 1. Each parameter set was simulated for 5 seconds and network activities were color-coded to indicate either synchronous network activity (‘sync’) or asynchronous network activity (‘async’). (C): Based on the criteria of (24), network simulations that showed a mean pairwise correlation (‘corr’) higher than 0.1 and a lower mean coefficient of variation (‘CV’) than 1 were labeled as synchronous (‘sync’). All other network simulations that did not meet these two criteria were labeled as asynchronous networks (‘async’).

**Fig. S2.**
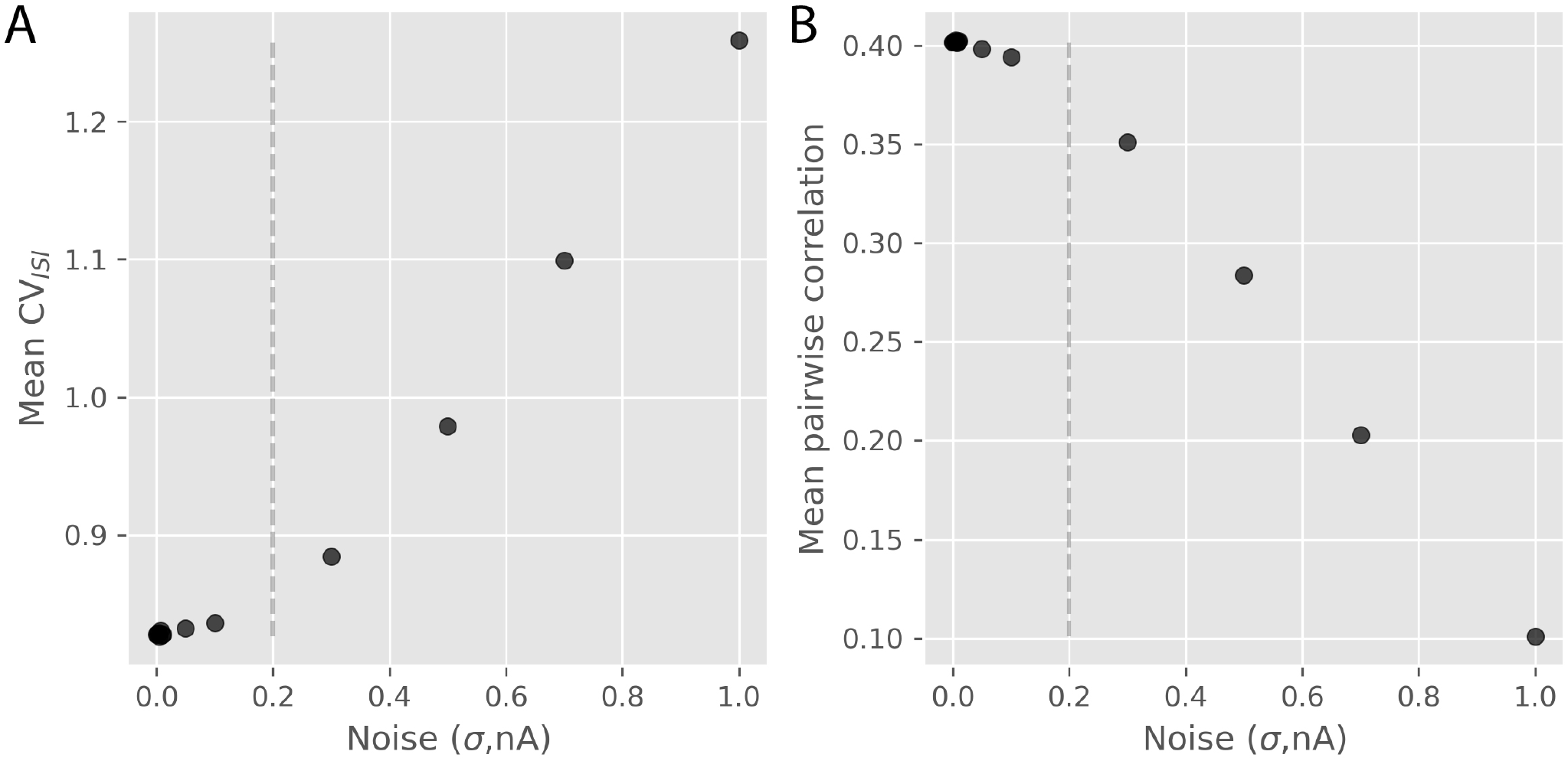
Noise-induced network activity changes in synchronous LIF simulations. (A): For noises that were below 0.2 nA (*≈* size of the leaky current), the mean CV_*ISI*_ of the simulated synchronous networks remained relatively stable. However, with larger noises, there was a rapid increase in mean CV_*ISI*_, and mean CV_*ISI*_ values larger than 1.0 were observed. (B): Similarly, with larger noise levels that exceed 0.2 nA, there was a steeper decrease in the mean pairwise correlation of the simulated synchronous networks compared to the case of lower noise levels.

**Fig. S3.**
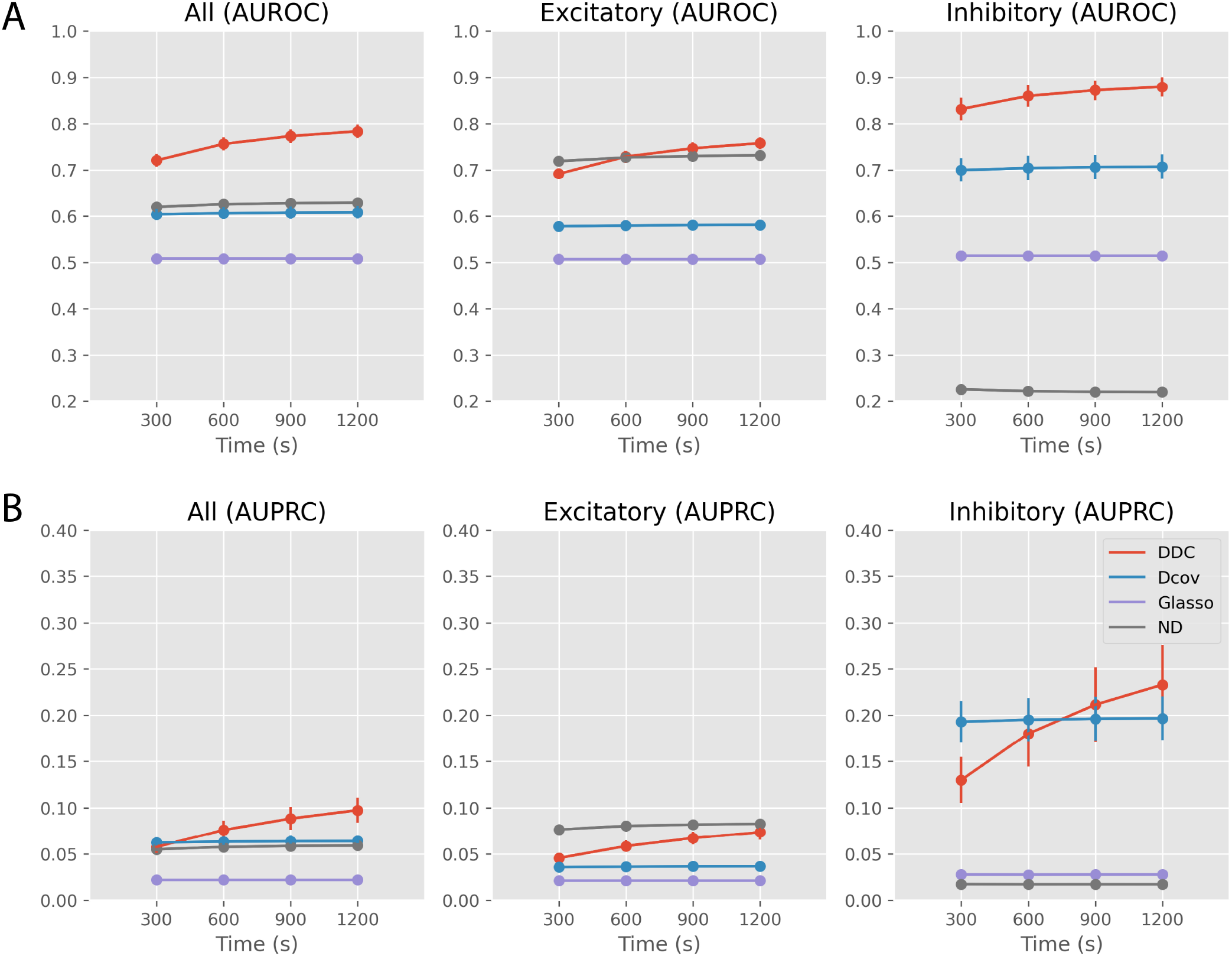
Inference performances of the multivariate methods (asynchronous networks). (A): AUROC values (mean *±* standard deviation) are presented for three evaluation conditions. (left): all ground truth connections, (middle): excitatory ground truth connections, (right): inhibitory ground truth connections. (B): AUPRC values are presented for three evaluation conditions. (left): all ground truth connections, (middle): excitatory ground truth connections, (right): inhibitory ground truth connections.

**Fig. S4.**
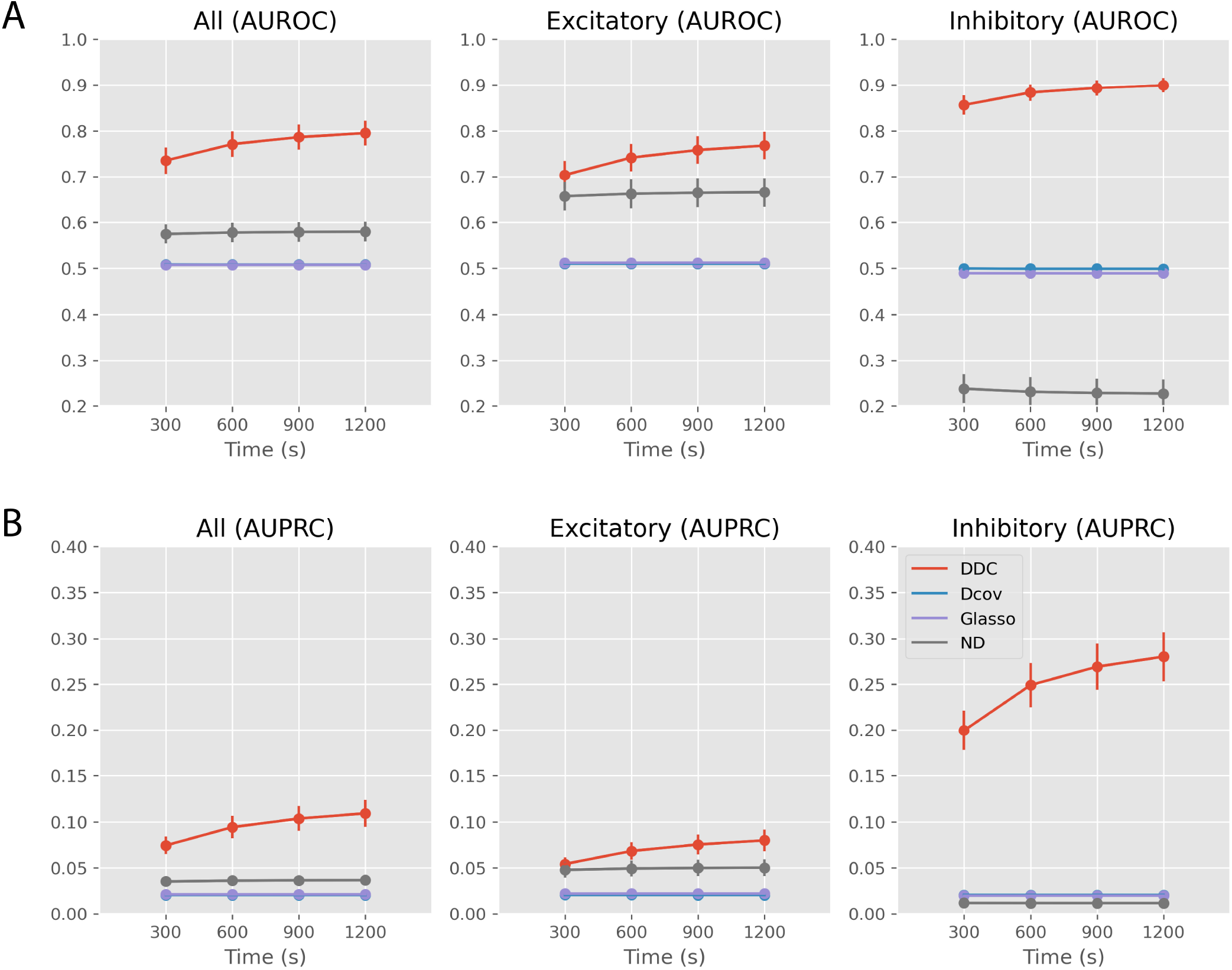
Inference performances of the multivariate methods (synchronous networks). (A): AUROC values (mean *±* standard deviation) are presented for three evaluation conditions. (left): all ground truth connections, (middle): excitatory ground truth connections, (right): inhibitory ground truth connections. (B): AUPRC values are presented for three evaluation conditions. (left): all ground truth connections, (middle): excitatory ground truth connections, (right): inhibitory ground truth connections.

## Footnotes

1 https://brian2.readthedocs.io/en/stable/

2 https://github.com/fabian-sp/GGLasso

3 https://fracdiff.github.io/fracdiff/

4 https://github.com/aestrivex/bctpy

## Notes

https://github.com/arahangua/FDDC

### Competing Interest Statement

The authors have declared no competing interest.

